# TDP-43 represses cryptic exon inclusion in FTD/ALS gene *UNC13A*

**DOI:** 10.1101/2021.04.02.438213

**Authors:** X. Rosa Ma, Mercedes Prudencio, Yuka Koike, Sarat C. Vatsavayai, Garam Kim, Fred Harbinski, Caitlin M. Rodriguez, H. Broder Schmidt, Beryl B. Cummings, David W. Wyatt, Katherine Kurylo, Georgiana Miller, Shila Mekhoubad, Nathan Sallee, Karen Jansen-West, Casey N. Cook, Sarah Pickles, Björn Oskarsson, Neill R. Graff-Radford, Bradley F. Boeve, David S. Knopman, Ronald C. Petersen, Dennis W. Dickson, Eric M. Green, William W. Seeley, Leonard Petrucelli, Aaron D. Gitler

**Author notes:** These authors contributed equally to this work (X. Rosa Ma, Mercedes Prudencio, Yuka Koike).

## Abstract

A hallmark pathological feature of neurodegenerative diseases amyotrophic lateral sclerosis (ALS) and frontotemporal dementia (FTD) is the depletion of RNA-binding protein TDP-43 from the nucleus of neurons in the brain and spinal cord. A major function of TDP-43 is as a repressor of cryptic exon inclusion during RNA splicing. Single nucleotide polymorphisms (SNPs) in *UNC13A* are among the strongest genome-wide association study (GWAS) hits associated with FTD/ALS in humans, but how those variants increase risk for disease is unknown. Here we show that TDP-43 represses a cryptic exon splicing event in *UNC13A*. Loss of TDP-43 from the nucleus in human brain, neuronal cell lines, and iPSC-derived motor neurons resulted in the inclusion of a cryptic exon in *UNC13A* mRNA and reduced UNC13A protein expression. Remarkably, the top variants associated with FTD/ALS risk in humans are located in the cryptic exon harboring intron itself and we show that they increase *UNC13A* cryptic exon splicing in the face of TDP-43 dysfunction. Together, our data provide a direct functional link between one of the strongest genetic risk factors for FTD/ALS (*UNC13A* genetic variants) and loss of TDP-43 function.

---

TDP-43, encoded by the *TARDBP* gene, is an abundant, ubiquitously expressed RNA-binding protein that normally localizes to the nucleus. It plays a role in fundamental RNA processing activities including RNA transcription, alternative splicing, and RNA transport ^1^. TDP-43 binds to thousands of pre-messenger RNA/mRNA targets ^2, 3^. Reduction in TDP-43 from an otherwise normal adult nervous system alters the splicing or expression levels of more than 1,500 RNAs, including long intron-containing transcripts ^2^. A major splicing regulatory function of TDP-43 is to repress the inclusion of cryptic exons during splicing ^4–7^. Unlike normal conserved exons, these cryptic exons lurk in introns and are normally excluded from mature mRNAs. When TDP-43 is depleted from cells, these cryptic exons get spliced into messenger RNAs, often introducing frame shifts and premature termination or even reduced RNA stability. However, the key cryptic splicing events that are integral to disease pathogenesis remain elusive.

*STMN2*, which encodes a regulator of microtubule stability called Stathmin-2, is the gene whose expression is most significantly reduced when TDP-43 is depleted from the human neuronal cell line SH-SY5Y and induced motor neurons ^8, 9^. *STMN2* harbors a cryptic exon (exon 2a) that is normally excluded from the mature *STMN2* mRNA. The first intron of *STMN2* contains a TDP-43 binding site. When TDP-43 is lost or its function is impaired, exon2a gets incorporated into the mature mRNA, suggesting that TDP-43 normally functions to repress the inclusion of exon 2a. Exon 2a harbors a stop codon and a polyadenylation signal – this results in truncated *STMN2* mRNA and 8-fold reduction of Stathmin-2 ^9^. Aberrant splicing and reduced Stathmin-2 levels seem to be a major feature of sporadic and familial ALS cases (except those with *SOD1* mutations) ^8, 9^ and in FTLD-TDP ^10^. The discovery of *STMN2* cryptic exon splicing in ALS and FTLD-TDP shines a bright light on one key mRNA target, but could there be others?

To discover cryptic splicing targets regulated by TDP-43 that may also play a role in disease pathogenesis, we utilized a recently generated RNA sequencing (RNA-Seq) dataset ^11^. To identify changes associated with loss of TDP-43 from the nucleus, the authors used fluorescence-activated cell sorting (FACS) to enrich neuronal nuclei with and without TDP-43 from postmortem FTD/ALS patient brain tissue and then performed RNA-seq to compare the transcriptomic profiles between TDP-43-positive and TDP-43-negative neuronal nuclei. They identified a multitude of interesting differentially expressed genes ^11^. Here, we re-analyzed the data to identify novel alternative splicing events impacted by the loss of nuclear TDP-43. We performed splicing analyses using two pipelines, MAJIQ ^12^ and LeafCutter ^13^, designed to detect novel splicing events (Fig. 1a). We identified 263 alternative splicing events (P(ΔΨ > 0.1) > 0.95)(ΔΨ, changes of local splicing variations between two conditions; P: probability) with MAJIQ and 152 with LeafCutter (P< 0.05). There were 65 alternatively spliced genes in common between both analyses (Fig. 1b), likely because each tool uses different definitions for transcript variations and different criteria to control for false positives. Among the alternatively spliced genes identified by both tools were *STMN2* and *POLDIP3*, both of which have been extensively validated as *bona fide* TDP-43 splicing targets ^8–10, 14^.

**Figure 1.**
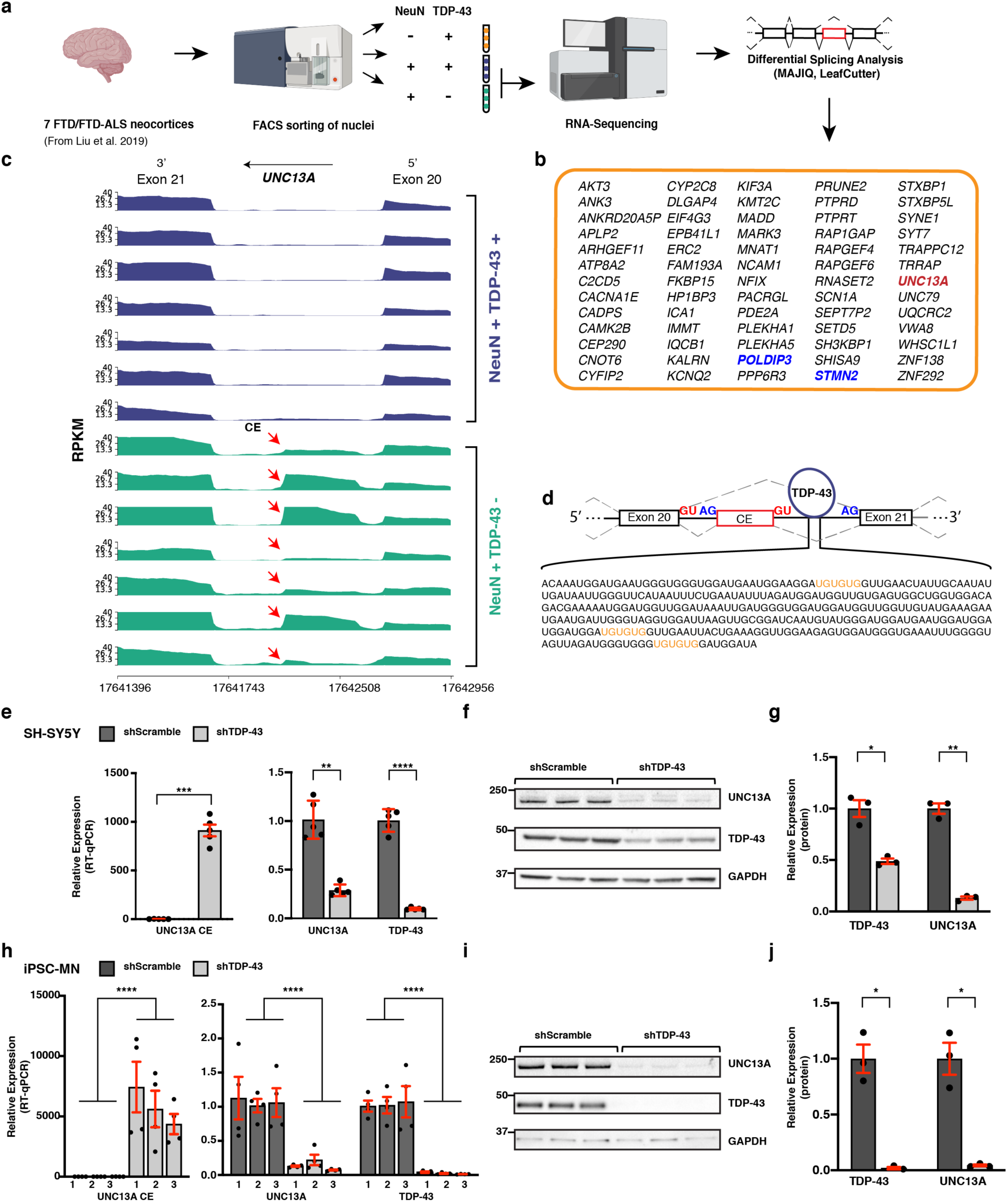
Nuclear depletion of TDP-43 causes cryptic exon inclusion in *UNC13A* RNA and reduced expression of UNC13A protein. **(a)** Splicing analyses were performed on RNA-sequencing results generated from TDP-43-positive and TDP-43-negative neuronal nuclei isolated from frontal cortices of seven FTD/FTD-ALS patients. FACS, fluorescence-activated cell sorting. **(b)** 65 alternatively spliced genes identified by both MAJIQ (P(ΔΨ > 0.1) > 0.95)(ΔΨ, changes of local splicing variations between two conditions) and LeafCutter (P < 0.05). **(c)** Visualization of RNA-sequencing alignment between exon 20 and exon 21 in *UNC13A* (hg38). Libraries were generated as described in (a). CE, cryptic exon. **(d)** iCLIP for TDP-43 (from Tollervey et al.^3^) indicates that TDP-43 binds to intron 20-21. An example of a region in intron 20-21 that is frequently bound by TDP-43 (shown by mapped reads). TDP-43 binding motif (UG)n is highlighted in orange. **(e and h)** RT-qPCR confirmed cryptic exon inclusion in *UNC13A* mRNA upon TDP-43 depletion in SH-SY5Y cells (5 independent cell culture experiments for each condition) (two sided-Welch Two Sample t-test, **P<0.01, ***P < 0.001, ****P<0.0001; mean ± s.e.m.) (e) and in 3 independent lines of iPSC derived motor neurons (iPSC-MNs) (4 independent cell culture experiments for each iPSC-MN) (h). *RPLP0* was used to normalize RT-qPCR. (Linear mixed model, ****P<0.0001; mean ± s.e.m.). **(f and i)** Immunoblotting for UNC13A and TDP-43 protein levels in SH-SY5Y cells (f) and iPSC-MNs (i) treated with scramble shRNA (shScramble) or TDP-43 shRNA (n=3). GAPDH served as a loading control. **(g and j)** Quantification of the blots in **(f and i),** respectively (two-sided Welch Two Sample t-test, *P<0.05, **P<0.01).

Unexpectedly, *UNC13A* was one of the genes with the most significant levels of alternative splicing in neurons with nuclear TDP-43 depletion (Fig. 1b). Depletion of TDP-43 resulted in the inclusion of a 128 bp cryptic exon between the canonical exons 20 and 21 (hg38; chr19: 17642414- 17642541) (Fig. 1 c and d, Extended Data Fig. 1; see Supplementary Note). This new exon (CE, for cryptic exon) was absent in wild type neuronal nuclei (Fig. 1c) and is not present in any of the known human isoforms of *UNC13A* ^15^. Furthermore, analysis of ultraviolet cross-linking and immunoprecipitation (iCLIP) data for TDP-43 ^3^ provides evidence that TDP-43 directly binds to the intron harboring this cryptic exon (shown by mapped reads) (Fig. 1d). Intron 20-21 of *UNC13A* and the CE sequence are conserved among most primates (Extended Data Fig. 2 a and b) but not conserved in mouse (Extended Data Fig. 2 c and d), similar to *STMN2* and other cryptic splicing targets of TDP-43 ^4, 8, 9^. Together, these results suggest that TDP-43 functions to repress the inclusion of a cryptic exon in the *UNC13A* mRNA transcript.

To determine whether TDP-43 directly regulates this *UNC13A* cryptic splicing event, we used shRNA to reduce TDP-43 levels in SH-SY5Y cells and quantitative reverse transcription PCR (RT-qPCR) to detect cryptic exon inclusion. The cryptic exon was present in cells with TDP-43 depletion but not in control shRNA-treated cells (Fig. 1e). Along with the increase in cryptic exon levels, there was a corresponding decrease in levels of the canonical *UNC13A* transcript upon TDP-43 depletion (Fig. 1e). By immunoblotting, we also observed a marked reduction in UNC13A protein in TDP-43-depleted cells (Fig. 1f). Reducing levels of TDP-43 in iPSC-derived motor neurons (iPSC-MNs) (Fig. 1 h to j; Extended Data Fig. 3 a and b) and excitatory neurons (i^3^Ns) derived from human iPS cells (Extended Data Fig. 4) also resulted in cryptic exon inclusion in *UNC13A* and a reduction in *UNC13A* mRNA and protein. We confirmed insertion of the 128 bp cryptic exon sequence into the mature transcript by direct sequencing of the RT-PCR product (* in Extended Data Fig. 3a). Thus, lowering levels of TDP-43 in human cells and neurons causes inclusion of a cryptic exon in the *UNC13A* transcript, resulting in decreased UNC13A protein.

*UNC13A* belongs to a family of genes originally discovered in *C. elegans* and was named based on the uncoordinated (*unc*) movements exhibited by animals with mutations in these genes ^16^, owing to deficits in neurotransmitter release. *UNC13A* encodes a large multidomain protein expressed in the nervous system, where it localizes to neuromuscular junctions and plays an essential role in the vesicle priming step, prior to synaptic vesicle fusion ^17–20^. Our *in vitro* studies demonstrate that the cryptic exon splicing event upon TDP-43 depletion caused marked reduction in UNC13A expression (Fig. 1 f and i). UNC13A is an essential neuronal protein because mice lacking Unc13a (also called Munc13-1) exhibit functional deficits at glutamatergic synapses, demonstrated by a lack of fusion-competent synaptic vesicles, and die within a few hours of birth ^19^. Our data suggest that depletion of TDP-43 leads to loss of this critical neuronal protein.

To extend our analysis of *UNC13A* cryptic exon (CE) inclusion to a larger collection of patient samples, we first analyzed a series of 117 frontal cortex brain samples from the Mayo Clinic Brain Bank and found a significant increase in *UNC13A* CE levels in FTLD-TDP patients compared to healthy controls (Fig. 2a and Extended Data Fig. 5a). Next, we analyzed brain samples from the New York Genome Center (NYGC). After quality control (Methods), we interrogated RNA-seq data from 1151 tissue samples from 413 individuals (with multiple tissues per individual), 330 of which are ALS or FTD patients. Because the FACS analysis by Liu et al. ^11^ indicates that neuronal nuclei with loss of TDP-43 represent only ∼7% of all neuronal nuclei and less than 2% of all cortical cells ^11^, we expected splicing analysis algorithms to struggle with detecting differentially spliced genes in RNA-seq data generated from bulk RNA sequencing. To overcome this problem, we specifically looked for reads that spanned the *UNC13A* exon 20-CE and CE-exon 21 junctions. Owing to noise generated from bulk sequencing, we scored the *UNC13A* splice variant as present if there were more than two reads spanning at least one of the exon-exon junctions. We identified 63 samples, from 49 patients, which met the above criteria. We detected *UNC13A* splice variants in nearly 50% of the frontal and temporal cortical tissues donated by patients with neuropathologically confirmed FTLD-TDP (Fig. 2b). We also detected the splice variants in some of the ALS patients whose pathology has not been confirmed (Extended Data Fig. 5b). Notably, we did not observe *UNC13A* CE in any of the samples from FTLD-FUS (n=9), FTLD-TAU (n=18) and ALS-SOD1 (n=22) patients, nor in any of the control samples (n=197) (Fig. 2b). Thus, *UNC13A* cryptic exon inclusion is a robust and specific facet of pathobiology in TDP-43 proteinopathies.

**Figure 2.**
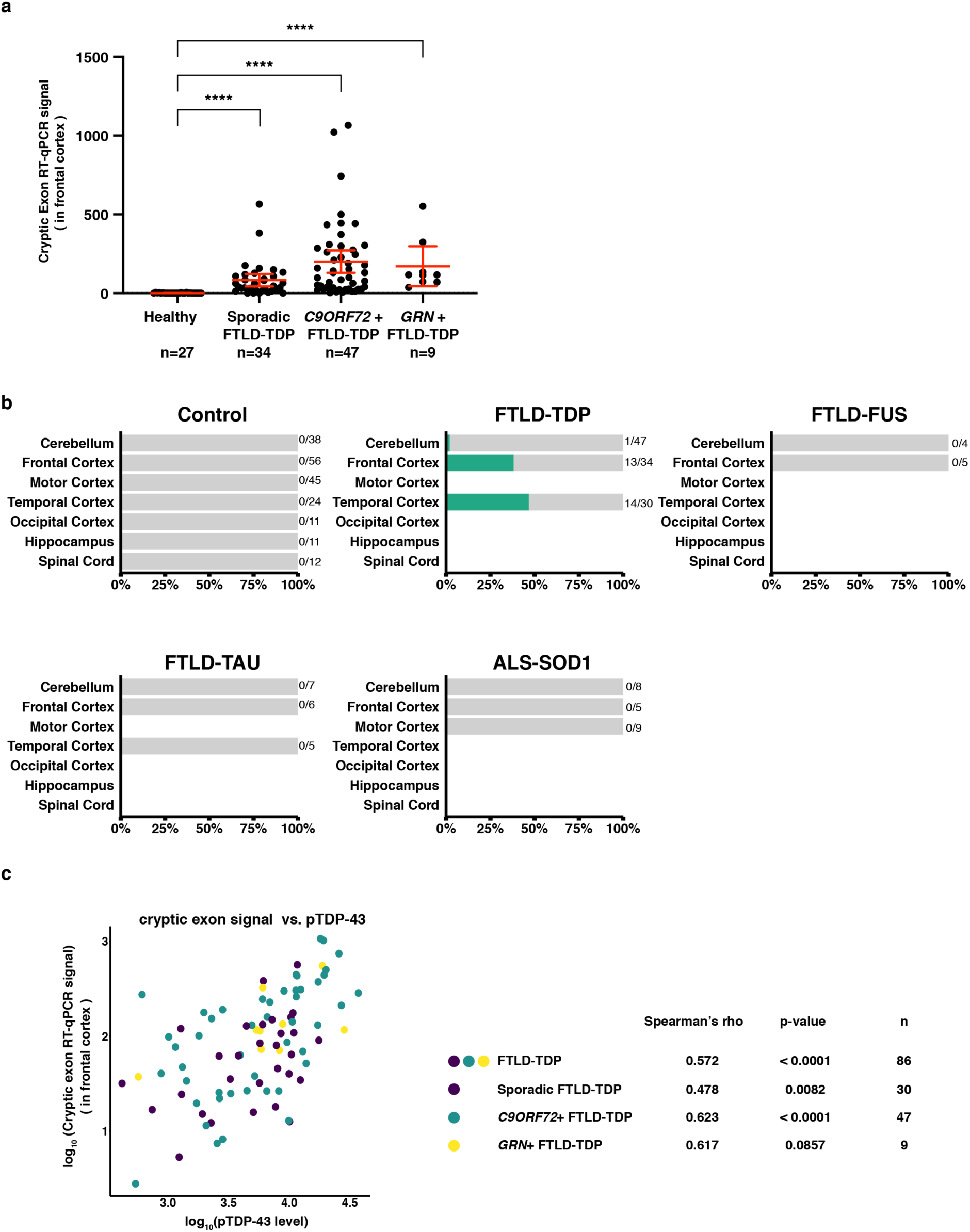
*UNC13A* cryptic exon inclusion in human TDP-43 proteinopathies. **(a)** *UNC13A* cryptic exon expression level is significantly increased in the frontal cortices of patients with FTLD-TDP pathologic diagnoses. *GAPDH* and *RPLP0* were used to normalize RT-qPCR (two- tailed Mann-Whitney test, ****P<0.0001; error bars represent 95% confidence intervals). **(b)** *UNC13A* cryptic exon is detected in nearly 50% of frontal cortical tissues and temporal cortical tissues from neuropathologically confirmed FTLD-TDP patients in bulk RNA-sequencing from the NYGC ALS Consortium cohort. The cryptic exon is also notably absent in tissues from healthy controls, FTLD-FUS, FTLD-TAU and ALS-SOD1 patients. **(c)** *UNC13A* cryptic exon signal is positively correlated with phosphorylated TDP-43 levels in frontal cortices of FTLD-TDP patients in Mayo Clinic Brain Bank (Spearman’s rho = 0.572, p-value <0.0001). Data points are colored according to patients’ reported genetic mutations. The correlation within each genetic mutation group and the corresponding p-value and sample size is also shown.

Once TDP-43 becomes depleted from the nucleus and accumulates in the cytoplasm, it becomes phosphorylated. Hyperphosphorylated TDP-43 (pTDP-43) is a key feature of pathology ^21^. To determine the relationship between pTDP-43 levels and *UNC13A* cryptic exon inclusion we analyzed data from 90 patients with FTLD-TDP in the Mayo Clinic Brain Bank for which we had RT-qPCR and pTDP-43 levels (Methods) from frontal cortices. We found a striking association between higher pTDP-43 levels and higher levels of *UNC13A* cryptic exon inclusion in patients with FTLD-TDP (Spearman’s rho = 0.572, P < 0.0001) (excluding 4 individuals who did not show *UNC13A* CE signal) (Fig. 2c and figure using raw data, Extended Data Fig. 5c). Thus, *UNC13A* cryptic exon inclusion is a common feature of multiple TDP-43 proteinopathies and to strongly correlate with the burden of TDP-43 pathology.

To visualize the *UNC13A* CE within single cells in the human brain, we designed custom BaseScope™ *in situ* hybridization probes that specifically bind to the exon 20-CE junction (Fig. 3a) or the exon 20-exon 21 junction (Extended Data Fig. 6). We designed the probes to span exon-exon junctions in order to minimize the possibility of binding to pre-mRNA. We used these probes for *in situ* hybridization, which was combined with immunofluorescence for NeuN (to detect neuronal nuclei) and TDP-43. We stained sections from the medial frontal pole of 4 FTLD-TDP patients and 3 controls (Fig. 3b and Extended Data Fig. 6a). In neurons showing loss of nuclear TDP-43 and accompanying cytoplasmic TDP-43 inclusions, we observed *UNC13A* CE-containing mRNA splice variants in the nucleus (Fig. 3b and Extended Data Fig. 6a). We observed 1-4 CE-containing mRNA puncta per nucleus. We did not observe puncta in the cytoplasm, perhaps since the cryptic exon introduces a premature stop codon, which could lead to non-sense mediated decay. Controls, however, had universally normal nuclear TDP-43 staining and showed no evidence of *UNC13A* cryptic splicing (Fig. 3c). Next, we sought to determine whether TDP-43 nuclear depletion is associated with reduced expression of canonical *UNC13A* mRNA. In control brain tissue, *UNC13A* mRNA was widely expressed in neurons across cortical layers (Extended Data Fig. 6 b and c). In patients, *UNC13A* mRNA was either absent or reduced in neurons showing TDP-43 pathology compared to neighboring neurons with normal nuclear TDP-43 (Extended Data Fig. 6c). These findings suggest that cryptic splicing of *UNC13A* is absent from controls and, in patients, is seen exclusively in neurons showing depleted nuclear TDP-43.

**Figure 3.**
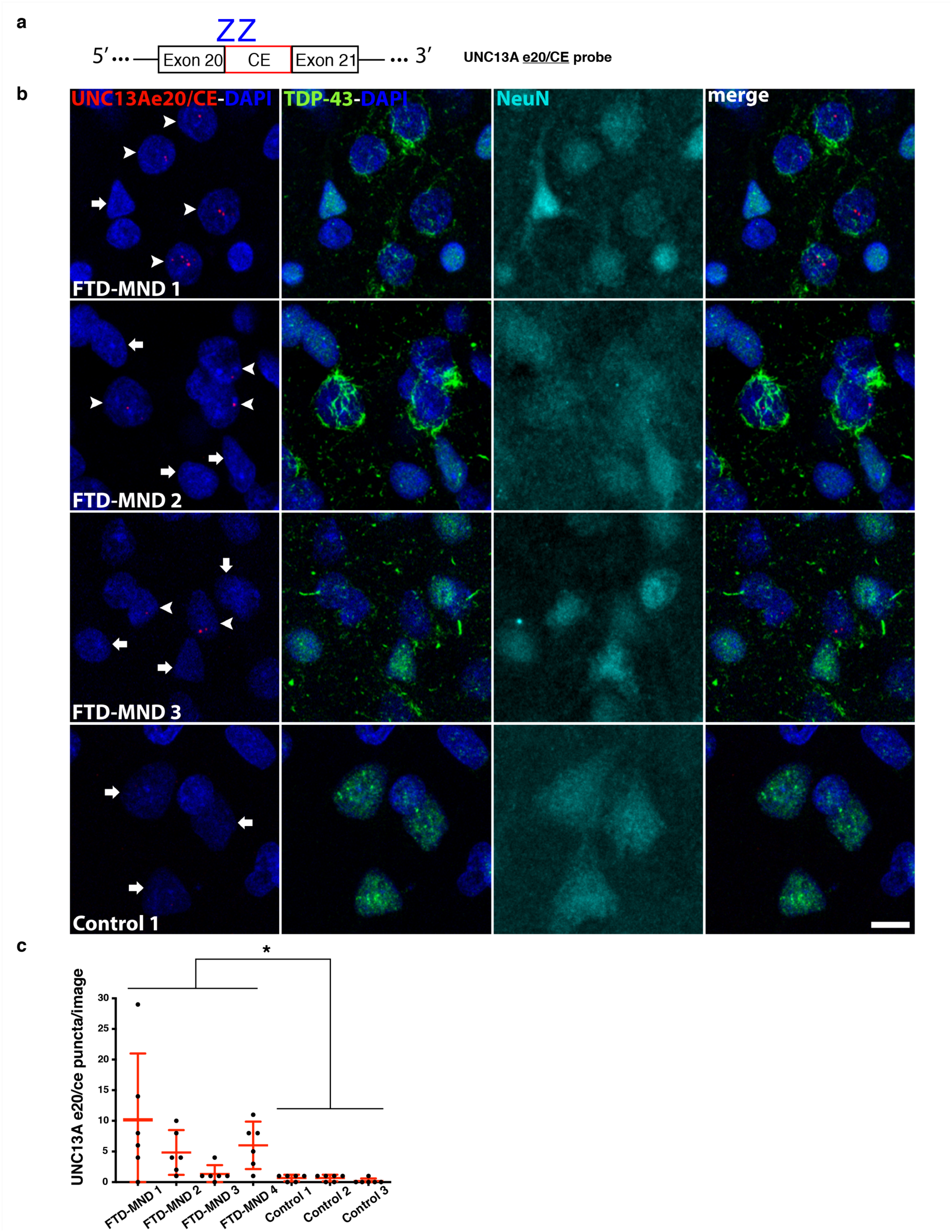
*UNC13A* cryptic splicing is associated with loss of nuclear TDP-43 in patients with FTD and motor neuron disease (MND). **(a)** The design of the UNC13A e20/CE BaseScope™ Z-stack images from layer 2-3 of medial frontal pole were captured, per subject, using a 63X oil objective and flattened into a maximum intensity projection image. Puncta counts per image were derived using the “analyze particle” plugin in ImageJ. Each data point represents the number of *UNC13A* cryptic exon puncta in a single image. The abundance of cryptic exons varies between patients but always exceeds the technical background of the assay, as observed in controls. Data are presented as mean +/- standard deviation (Linear mixed model, *P<0.05).

*UNC13A* is one of the top GWAS hits for ALS and FTD-ALS, replicated across multiple studies ^22–27^. SNPs in *UNC13A* are associated with increased risk of sporadic ALS ^23^ and sporadic FTLD- TDP pathology, especially Type B, the subtype associated with FTD-ALS ^22^. In addition to increasing susceptibility to ALS, SNPs in *UNC13A* are associated with shorter survival in ALS patients ^28–31^. But the mechanism by which genetic variation in *UNC13A* increases risk for ALS and FTD is unknown. Remarkably, the two most significantly associated SNPs, rs12608932 (A>C) and rs12973192 (C>G), are both located in the same intron that we found harbors the cryptic exon, with rs12973192 located right in the cryptic exon itself (Fig. 4a). This immediately suggested the hypothesis that these SNPs (or other genetic variation nearby tagged by these SNPs) might make *UNC13A* more vulnerable to cryptic exon inclusion upon TDP-43 depletion. To test this hypothesis, we analyzed the percentage of RNA-seq reads (Extended Data Fig. 7 a and b) that mapped to intron 20-21 that support the inclusion of the cryptic exon. Among the seven patients included in the initial splicing analysis (Fig. 1a), 2 out of 3 who were homozygous (G/G) and the one patient that was heterozygous (C/G) for the risk allele at rs12973192 showed inclusion of the cryptic exon in almost every *UNC13A* mRNA that was mapped to intron 20-21. In contrast, the 3 other patients who were homozygous for the reference allele (C/C) showed much less inclusion of the cryptic exon (Extended Data Fig. 7 a and b). Another way to directly assess the impact of the *UNC13A* risk alleles on cryptic exon inclusion is to measure allele imbalance in RNAs from individuals who are heterozygous for the risk allele. In other words, is there an equal number of RNA transcripts with CE inclusion produced from the risk allele as from the reference allele? Or are there more from the risk allele? Two of the iPSC-MN lines that we used to detect cryptic exon inclusion upon TDP-43 knockdown (Fig. 1h, iPSC-MN1 and iPSC-MN3) are heterozygous (C/G) at rs12973192. We sequenced the RT-PCR product that spans the cryptic exon and analyzed the allele distribution from these two samples as well as the one patient sample from the original RNA-seq dataset (Fig. 1a) that is heterozygous (C/G) at rs12973192 (Extended Data Fig. 7b). We found a significant difference between the percentage of C and G alleles in the spliced variant, with higher inclusion of the risk allele (Fig. 4b and Extended Data Fig. 7c).

**Figure 4.**
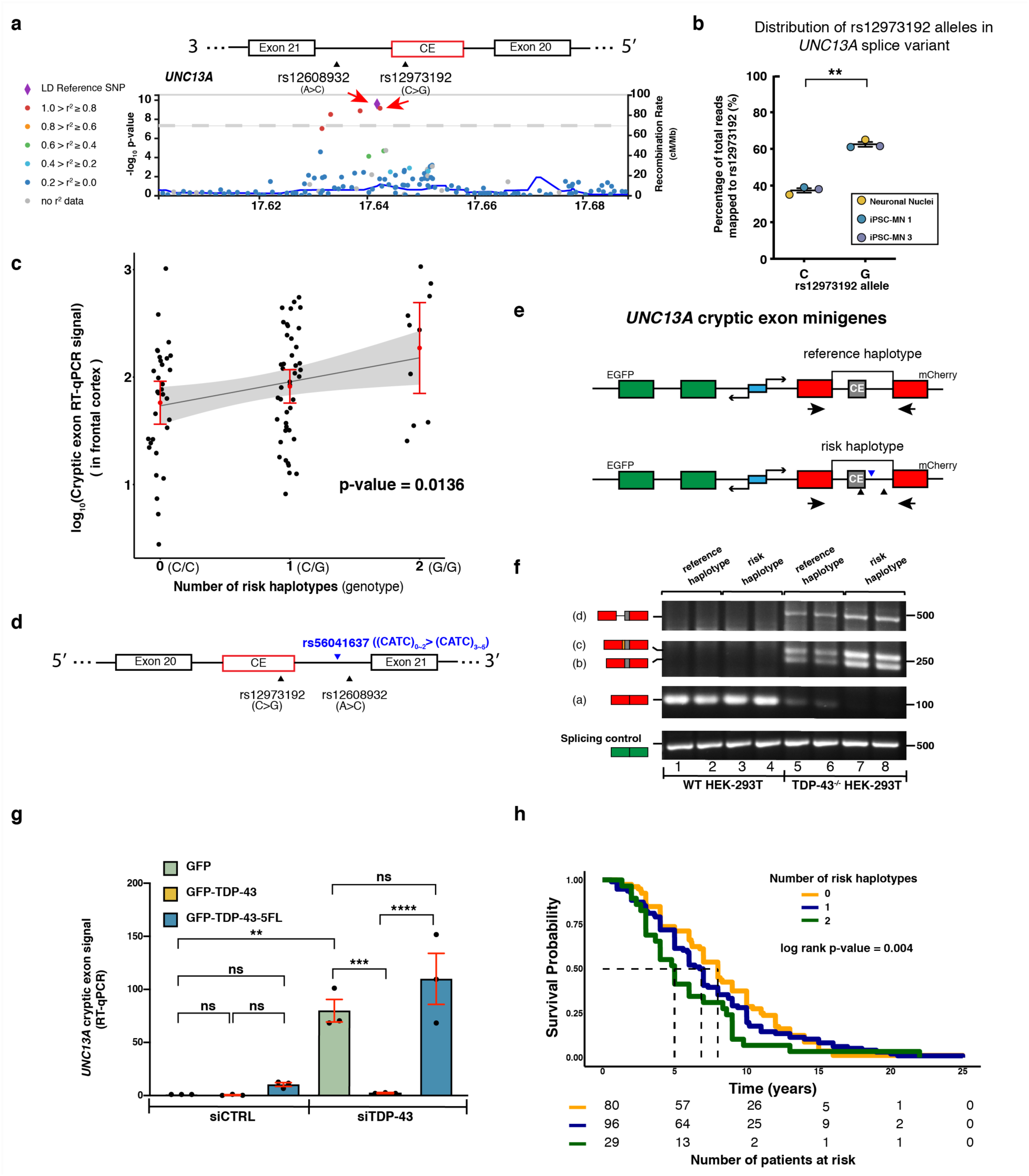
*UNC13A* risk haplotype associated with ALS/FTD susceptibility potentiates cryptic exon inclusion when TDP-43 is dysfunctional. (a) LocusZoom plot showing SNPs associated with ALS/FTD in *UNC13A*. rs12608932, the most significant GWAS hit is chosen to be the reference. Other SNPs are colored based on their levels of linkage equilibrium with rs12608932 in EUR population. The two SNPs in intron 20-21 (black triangles), rs12608932 and rs12973192 are in strong linkage disequilibrium. **(b)** There is a higher percentage inclusion of the risk allele (G) at rs12973192 in *UNC13A* splice variant (two-sided paired t-test, **P = 0.0094). **(c)** Linear regression model shows a strong correlation between the abundance of *UNC13A* cryptic exon and the number of risk haplotypes. Normality of residuals is tested by Shapiro-Wilk normality test (p-value = 0.2604). **(d)** Diagram of the location of rs56041637 relative to the two known ALS and FTD-ALS GWAS hits and *UNC13A* cryptic exon. **(e)** Design of *UNC13A* cryptic exon minigene reporter constructs and the location of the primer pair used for RT-PCR. Transcription of GFP and mCherry is controlled by a bidirectional promoter (blue). Black triangles represent the locations of genetic variants as shown in (d). **(f)** Splicing of the minigene reporters was assessed in WT and TDP-43^-/-^ HEK293T cells. HEK293T cells do not endogenously express *UNC13A*. The PCR products represented by each band are marked to the left of each gel. In addition to the inclusion of cryptic exon (b), some splice variants showed inclusion of the other two cryptic splicing products (c and d) (Extended Data Fig. 1a and see Supplementary Note). The risk haplotype- carrying minigene showed an almost complete loss of canonical splicing product (a) and an increase in alternatively spliced products (b, c, and d). **(g)** In HeLa cells expressing a different *UNC13A* minigene reporter (Extended Data Fig. 8a), depletion of TDP-43 by siRNA, resulted in inclusion of the cryptic exon, which was rescued by expressing an siRNA-resistant form of TDP- 43 (GFP-TDP-43) but not by an RNA-binding deficient mutant TDP-43 (GFP-TDP-43-5FL) (One-way ANOVA with Sidak’s multiple comparisons test, ns, P > 0.05, ***P < 0.001, ****P<0.0001). **(h)** Survival curves of FTLD-TDP patients stratified based on the number of risk haplotypes they carry (0, 1, or 2). Patients who are heterozygous and homozygous for the risk haplotype had shorter survival time after disease onset (n=205, Mayo Clinic Brain Bank) (Score (logrank) test, p-value = 0.004). Dashed lines mark the median survival for each genotype. The effect of the risk haplotype is modeled as an additive model using Cox multivariable analysis adjusted for genetic mutations, sex and age at onset. The risk table is shown at the bottom. Summary results of the analysis are in Extended Data Fig. 9a.

Given this evidence for an effect of the risk allele on cryptic exon inclusion, we extended our analysis by genotyping FTLD-TDP patients with cryptic exon signal (n = 86) in the Mayo Clinic Brain Bank dataset for the *UNC13A* risk alleles at rs12973192 and rs12608932. Because these two SNPs are in high linkage disequilibrium in the European population ^32^, we consider them to be on the same haplotype. Thus, we refer to the haplotype that contains reference alleles at both SNPs as the reference haplotype, and the haplotype that contains risks alleles as the risk haplotype. We excluded the one patient who is homozygous for the reference allele (C/C) at rs12973192 but heterozygous (A/C) at rs12608932. The rest of the patients (n=85) have exactly the same number of risk alleles at both loci, indicating that it’s very likely the patients are carriers of the reference haplotype or the risk haplotype. Using a linear regression model, we found a strong correlation between the number of risk haplotypes and the abundance of *UNC13A* cryptic exon inclusion (Fig. 4c and figure using raw data, Extended Data Fig. 7d). Taken together, these data suggest that genetic variation in *UNC13A* that increases risk for ALS and FTD in humans promote cryptic exon inclusion upon TDP-43 nuclear depletion.

GWAS SNPs typically do not cause the trait but rather “tag” other neighboring genetic variation^33^. Thus, a major challenge in human genetics is to go from a GWAS hit to identifying the causative genetic variation that increases risk for disease ^34^. A LocusZoom ^35^ plot (Fig. 4a) generated using results from an ALS GWAS ^36^ suggests that the strongest association signal on *UNC13A* is indeed in the region surrounding the two lead SNPs (rs12973192 and rs12608932). To look for other genetic variants in intron 20-21 that might also cause risk for disease by influencing cryptic exon inclusion but were not included in the original GWAS, we analyzed genetic variants identified in whole genome sequencing data of 297 ALS patients of European descent (Answer ALS). We searched for novel genetic variants that could be tagged by the two SNPs by looking for other loci in intron 20-21 that are in linkage disequilibrium with both rs12608932 and rs12973192. We found one that fit these criteria: rs56041637 (Extended Data Fig. 7e). rs56041637 is a CATC-repeat insertion. In the patient dataset, we observed that patients who are homozygous for the risk alleles at both rs12608932 and rs12973192 tend to have 3 to 5 CATC-repeats at rs56041637; patients who are homozygous for reference alleles at both rs12608932 and rs12973192 tend to have shorter (0 to 2) repeats at rs56041637 (Fig. 4d). Thus, in addition to the two lead GWAS SNPs (rs12608932 and rs12973192), we now nominate rs56041637, as potentially contributing to risk for disease by making *UNC13A* more vulnerable to cryptic exon inclusion when TDP-43 is depleted from the nucleus.

To directly test if these three variants in *UNC13A* — which are part of the risk haplotype — increase cryptic exon inclusion upon TDP-43 depletion, we synthesized minigene reporter constructs (Fig. 4e). The reporter uses a bidirectional promoter to co-express EGFP containing a canonical intron and mCherry that is interrupted by *UNC13A* intron 20-21 from either the reference haplotype or the risk haplotype. We transfected WT and TDP-43-knock-out (TDP43^−/−^) HEK- 293T cells ^37^, which do not express *UNC13A* endogenously, with each minigene reporter construct. Using RT-PCR, we found that both versions of intron 20-21 were efficiently spliced out in WT cells (Fig. 4f, lane 1-4). However, in TDP43^−/−^ cells there was a decrease in completely intron-free splicing products and a concomitant increase in cryptic splicing products (Fig. 4f, lane 5-6). Strikingly, in TDP-43^−/−^ cells transfected with the minigene construct harboring the risk haplotype in the intron, there was an even greater decrease in complete intron 20-21 splicing, and a proportional increase in cryptic splicing products (Fig. 4f, lane 7-8; see Supplementary Note). The transcript levels of the EGFP control remained constant across all conditions, verifying equal reporter expression levels and the integrity of the splicing machinery independent of TDP-43. To corroborate our findings, we tested a different minigene reporter construct (Extended Data Fig. 8a) in TDP-43 deficient HeLa cells, which also resulted in splicing defects (Fig. 4g). Expression of shRNA-insensitive WT TDP-43 rescued these splicing defects, while an RNA binding-deficient mutant with five phenylalanine residues mutated to leucine (5FL) failed to do so (Fig. 4g and Extended Data Fig. 8b). Together, these results provide direct functional evidence that 1) TDP-43 regulates splicing of *UNC13A* intron 20-21 and 2) genetic variants associated with ALS and FTD susceptibility in humans potentiate cryptic exon inclusion when TDP-43 is dysfunctional.

To examine whether these SNPs affect survival in FTLD-TDP patients (n=205) in the Mayo Clinic Brain Bank, we evaluated the association of the risk haplotype with survival time after disease onset. Using Cox multivariable analysis adjusting for other factors (genetic mutations, sex, age at onset) known to influence survival, the risk haplotype was associated with survival time under an additive model (Fig. 4h). The number of risk haplotypes an individual carries was a strong prognostic factor (Extended Data Fig. 9a). The association remained significant under a dominant model (Extended Data Fig. 9 b and c) and a recessive model (Extended Data Fig. 9 d and e), indicating that carrying the risk haplotype reduces patient survival time after disease onset, consistent with previous analyses ^28–31^. Thus, genetic variants in *UNC13A* that increase cryptic exon inclusion are associated with decreased survival in patients.

Here, we found that TDP-43 regulates a cryptic splicing event in the FTD/ALS risk gene *UNC13A*. The most significant genetic variants associated with disease risk, including a novel variant nominated here, are located within the intron harboring the cryptic exon itself. Brain samples from FTLD-TDP patients carrying these SNPs exhibited more *UNC13A* cryptic exon inclusion than those from FTLD-TDP patients lacking the risk alleles. These risk alleles seem insufficient to cause cryptic exon inclusion because the cryptic exon is not detected in RNA-seq data from healthy control samples (GTEx) ^15^ and our functional studies indicate that TDP-43 dysfunction is required for *UNC13A* cryptic exon inclusion. Instead, the *UNC13A* risk alleles exert a TDP-43 loss-of-function-dependent disease modifying effect. We propose that the *UNC13A* risk alleles might act as a kind of Achilles’ heel – lurking under the surface, not causing problems until TDP-43 becomes dysfunctional. We do not expect to observe severe loss-of-function mutations in the *UNC13A* coding region because these would result in early lethality, as seen in mice ^19^. The discovery of a novel TDP-43-dependent cryptic splicing event in a *bona fide* FTD-ALS risk gene opens up a multitude of new directions for validating *UNC13A* as a biomarker and therapeutic target in ALS and FTD. It still remains a mystery why TDP-43 pathology is associated with ALS, FTLD-TDP, or even other aging-related neuropathological changes ^38^. TDP-43 dysfunction-related cryptic splicing plays out across the diverse regional and neuronal landscape of the human brain. It is tempting to speculate that in addition to *STMN2*, and now *UNC13A,* there could be specific portfolios of other important cryptic exon splicing events (and genetic variations that increase or decrease susceptibility to some of these events) that contribute to heterogeneity in clinical manifestation of TDP-43 dysfunction.

## Acknowledgments

This work was supported by NIH grants R35NS097263(10) (A.D.G.), R35NS097273(17) (L.P.), R01NS104437 (W.W.S.), F32 NS116208-02 (C.M.R), the Robert Packard Center for ALS Research at Johns Hopkins (L.P., A.D.G.), P30AG06267 (R.C.P.), U01AG006786 (R.C.P.), 2T32HG000044-21 NIHGRI training grant (X.R.M.), and the Brain Rejuvenation Project of the Stanford Neurosciences Institute (A.D.G.). G.K. is supported by a fellowship from the Stanford Knight-Hennessy Scholars Program. The UCSF Neurodegenerative Disease Brain Bank receives funding support from NIH grants P30AG062422, P01AG019724, U01AG057195, and U19AG063911, as well as the Rainwater Charitable Foundation and the Bluefield Project to Cure FTD. Some of the computing for this project was performed on the Sherlock cluster. We would like to thank Stanford University and the Stanford Research Computing Center for providing computational resources and support that contributed to these research results.

## Data availability

All data used in this study are available upon request.

## Code availability

**All codes** used in this study are available upon request.

## Supplementary Note

From the splicing analysis using the RNA-Seq data from Liu et al (Fig. 1a), we found that depletion of TDP-43 introduces two alternative 3’ splice acceptor sites in intron 20-21: one is near chr19:17642591(ΔΨ=0.05184) and the other one is at chr19:17642541(ΔΨ=0.48865). They share the same alternative 5’ splice donor site near chr19:17642414 (ΔΨ=0.772) (Extended Data Fig. 1a). This creates two different cryptic exons. Since we saw much higher usage of the chr19:17642541 3’ splicing acceptor, we focus on the 128 bp cryptic exon (CE) defined by this 3’ splice acceptor and the alternative 5’ splice acceptor. The other version of the cryptic exon (hg38; chr19: 17642414-17642591) is 50 bp longer than the more abundant version (CE-2). RT-PCR amplifying the exon 20-exon 21 region (Extended Data Fig. 3) indicates that there is also a version of the cryptic exon which uses the same alternative 5’ splice donor but includes the entire intron between exon 20 and CE (CE-3) (hg38; chr19: 17642844-17642414). We did not observe CE-2 on the gel because the two are very close in size. It is likely that CE-3 was not observed in the splicing analysis (Fig. 1a) because of the stringent parameter choice. All three cryptic exons were detected in the mini-gene experiment (Fig. 4h). CE-2 is 178 bp and CE-3 is 431bp, indicating that, similar to CE, both CE-2 and CE-3 would introduce a pre-mature stop codon and lead to dysregulation of UNC13A protein expression. The primer pair we used for RT-qPCR for this study spans the CE-exon 21 junction, which is the same as CE-2-exon 21 and CE-3-exon 21 junctions. Therefore, the primer pair can detect all three versions of the cryptic exon.

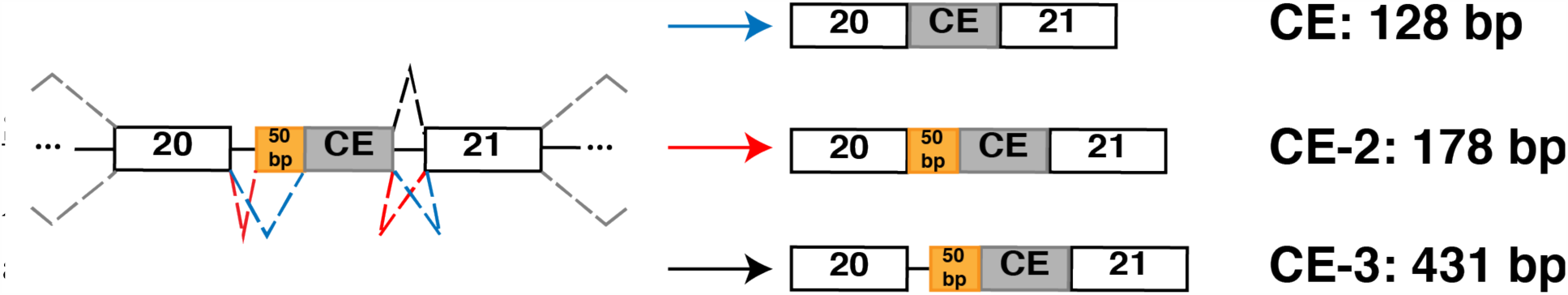

**Extended Data Fig. 1.**
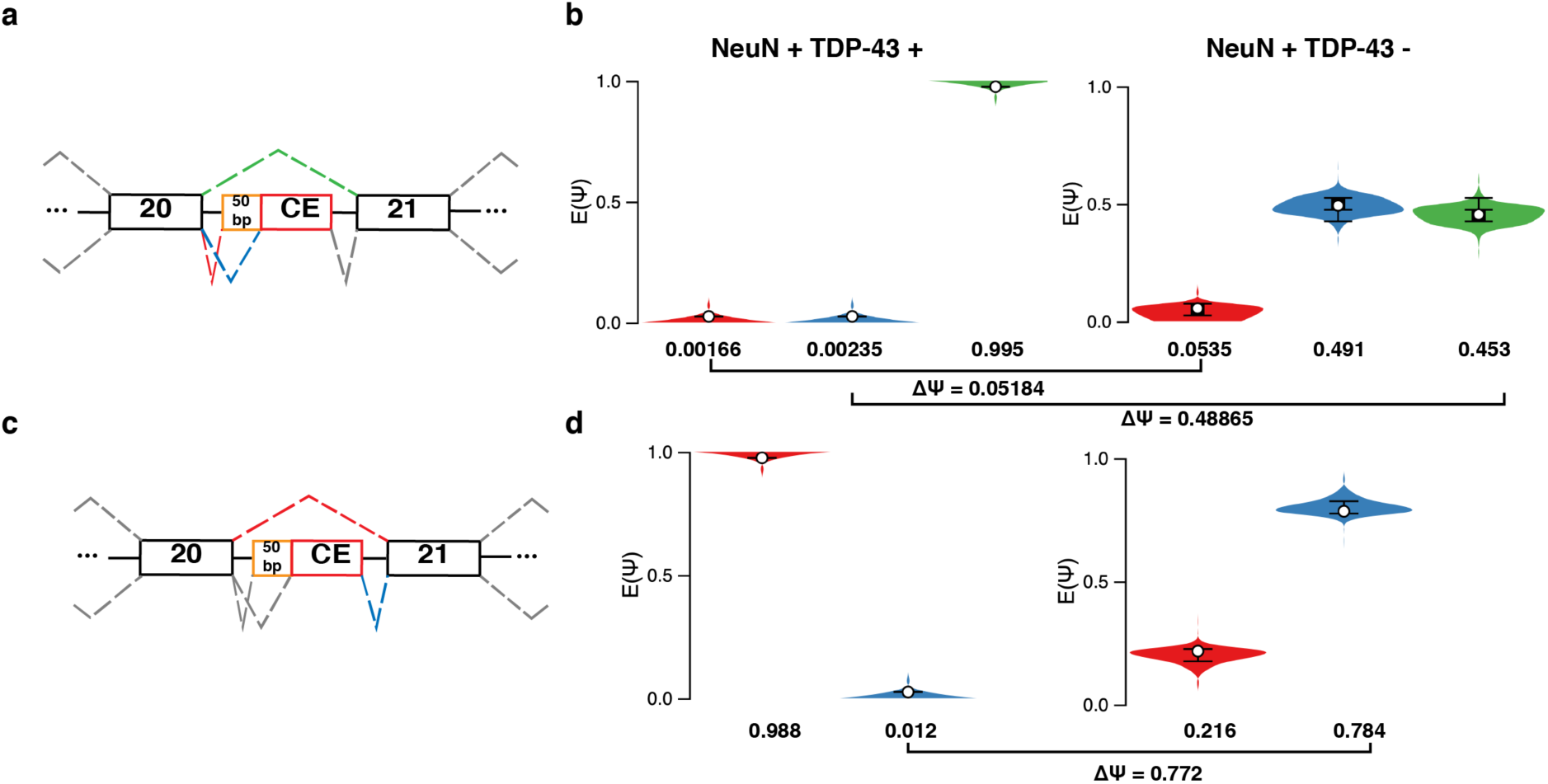
Splicing analysis using MAJIQ demonstrates inclusion of cryptic exon between exon 20 and exon 21 of *UNC13A*. **(a and b)** Depletion of TDP-43 introduces two alternative 3’ splicing acceptors in the intron 20-21: one is near chr19:17642591(ΔΨ=0.05184) and the other one is near chr19:17642541(ΔΨ=0.48865). (c and d) An alternative 5’ splicing donor is also introduced near chr19:17642414 (ΔΨ=0.772). Since we saw much higher usage of the chr19:17642541 3’ splicing acceptor (b), the focus of the paper is on the 128 bp cryptic exon defined by this 3’ splicing acceptor and the alternative 5’ splicing acceptor **(c)**. **(a and c)** Splice graphs showing the inclusion of the cryptic exon (CE) between exon 20 and exon 21 of *UNC13A*. **(b and d)** Violin plots corresponding to (a and c) respectively. Each violin in (b and d) represents the posterior probability distribution of the expected relative inclusion (PSI or Ψ) for the color matching junction in the splice graph. The tails of each violin represent the 10th and 90th percentile. The box represents the interquartile range with the line in the middle indicating the median. The white circles mark the expected PSI (E[Ψ]). The change in the relative inclusion level of each junction between two conditions is referred to as ΔΨ or ΔPSI^12^.

**Extended Data Fig. 2.**
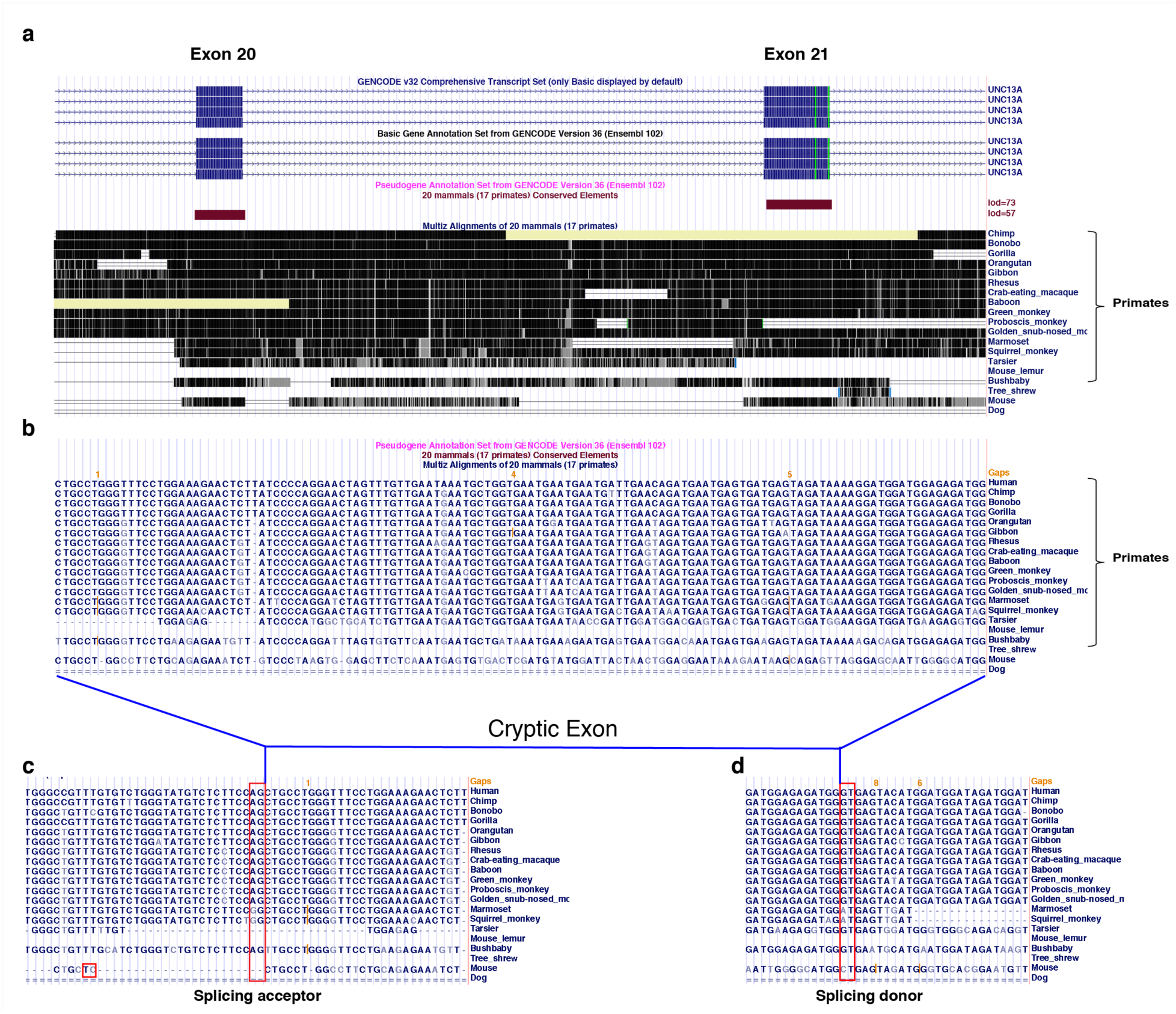
Intron 20-21 of *UNC13A* is conserved among most primates. The Primates Multiz Alignment & Conservation track on UCSC ^39^ genome browser (http://genome.ucsc.edu) includes 20 mammals, 17 of which are primates. **(a)** Exon 20 and exon 21 of *UNC13A* is well conserved among mammals. However, intron 20-21 **(a)**, the cryptic exon **(b)**, and the splicing acceptor site upstream of the cryptic exon **(c)** and splicing donor site downstream of the cryptic exon **(d)** are only conserved in primates.

**Extended Data Fig. 3.**
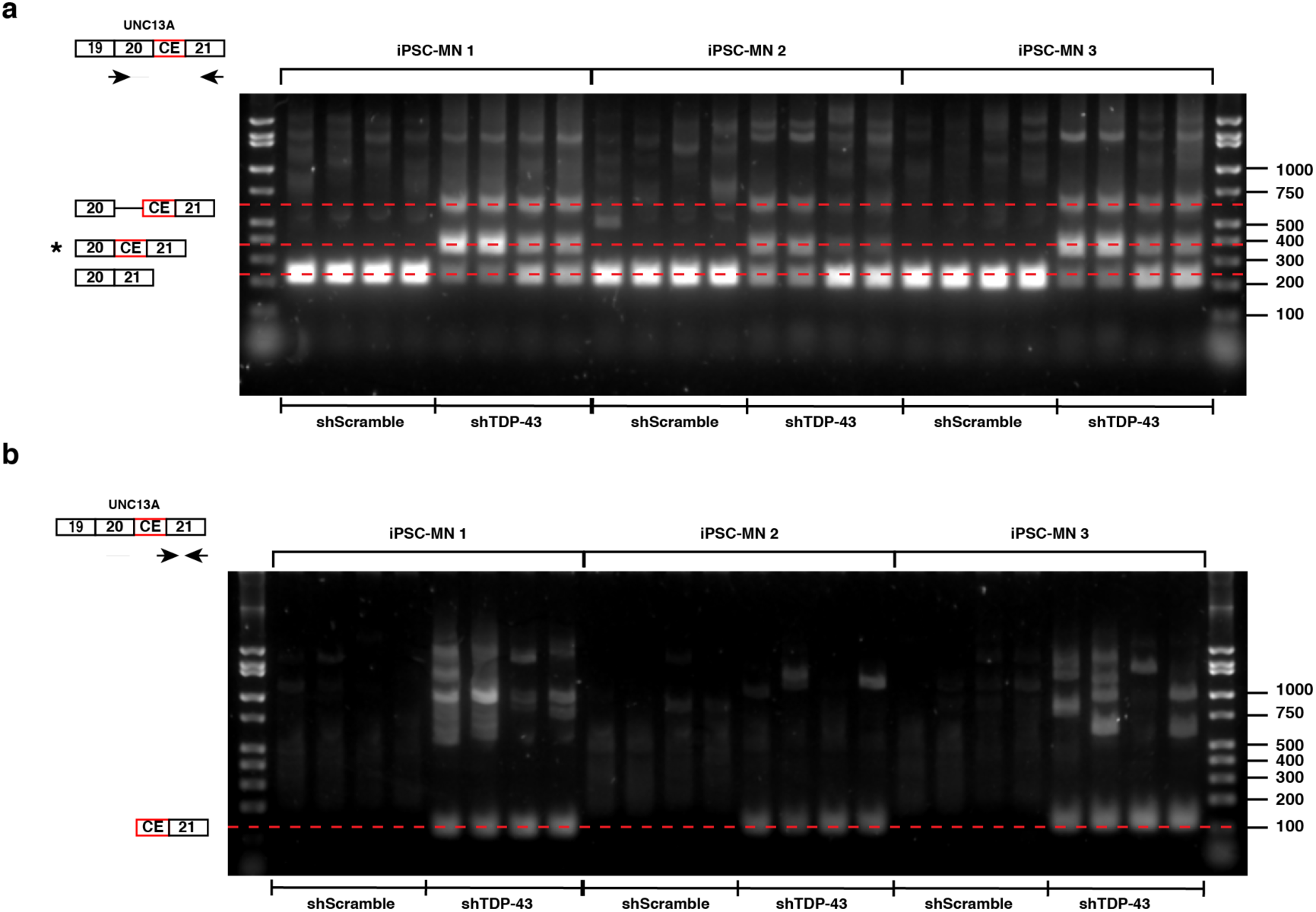
Depletion of TDP-43 from iPSC derived motor neurons (iPSC-MNs) leads to cryptic exon inclusion in *UNC13A*. **(a)** RT-PCR confirmed the expression of the cryptic exon-containing *UNC13A* splice variant upon TDP-43 depletion in three independent iPSC-MNs (4 independent cell culture experiments for each iPSC-MN and condition). In addition to the splice variant containing the cryptic exon, we also detected inclusion of the complete intron upstream of the cryptic exon (Fig. 4g, see Supplementary Note). The PCR products represented by each band are marked to the left of each gel. The location of the PCR primer pair used is shown on top of each gel image. **(b)** The PCR primer pairs spanning the cryptic exon and exon 21 junction confirms cryptic exon inclusion only occurs upon TDP-43 knockdown.

**Extended Data Fig. 4.**
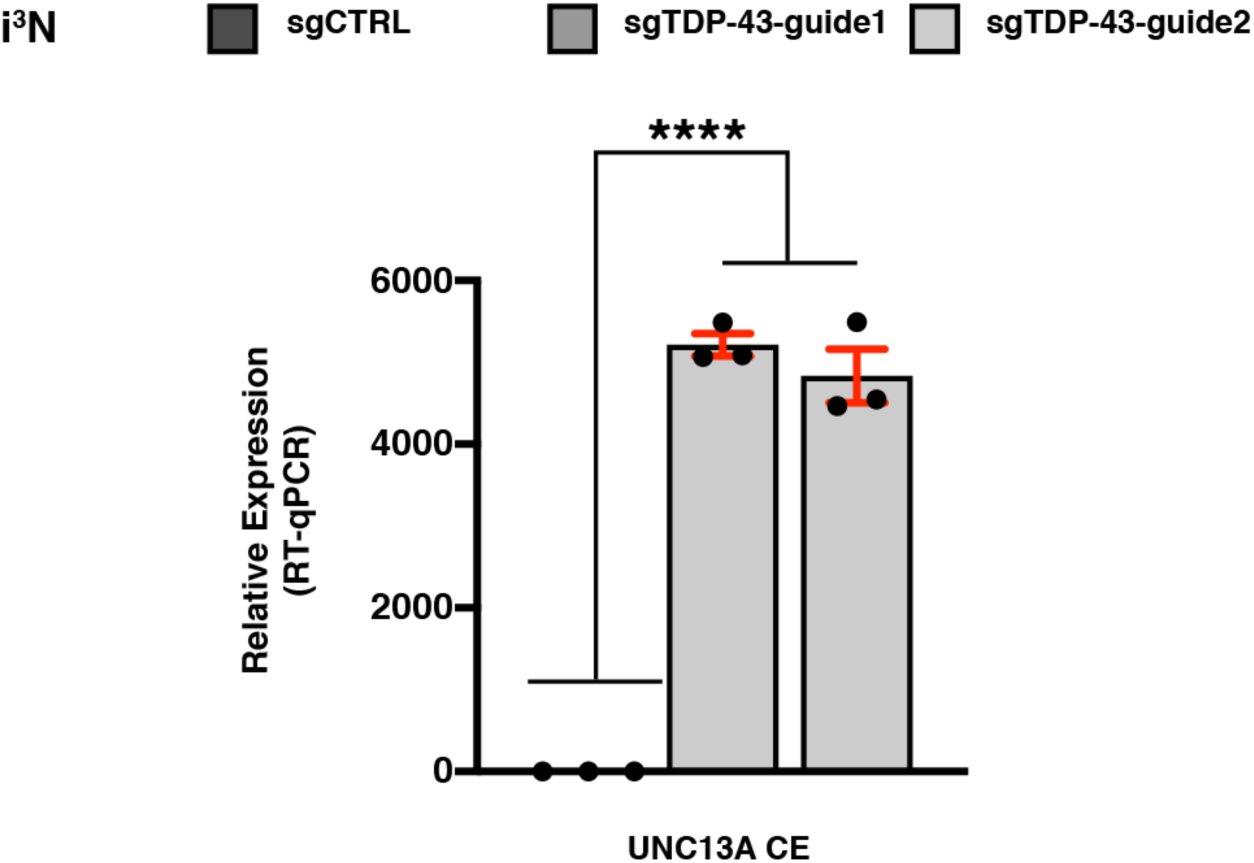
Depletion of TDP-43 causes *UNC13A* cryptic exon inclusion in i^3^Ns. RT-qPCR analyses confirmed the inclusion of *UNC13A* cryptic exon upon TDP-43 depletion in neurons derived from human iPS cells (i^3^Ns). TDP-43 was depleted by expressing two different sgRNAs: sgTDP-43-guide1 and sgTDP-43-guide2 in i^3^Ns stably expressing CRISPR inactivation machinery (CRISPRi). *RPLP0* and *GAPDH* were used to normalize RT-qPCR (Linear mixed model, ****P<0.0001; mean ± s.e.m.).

**Extended Data Fig. 5.**
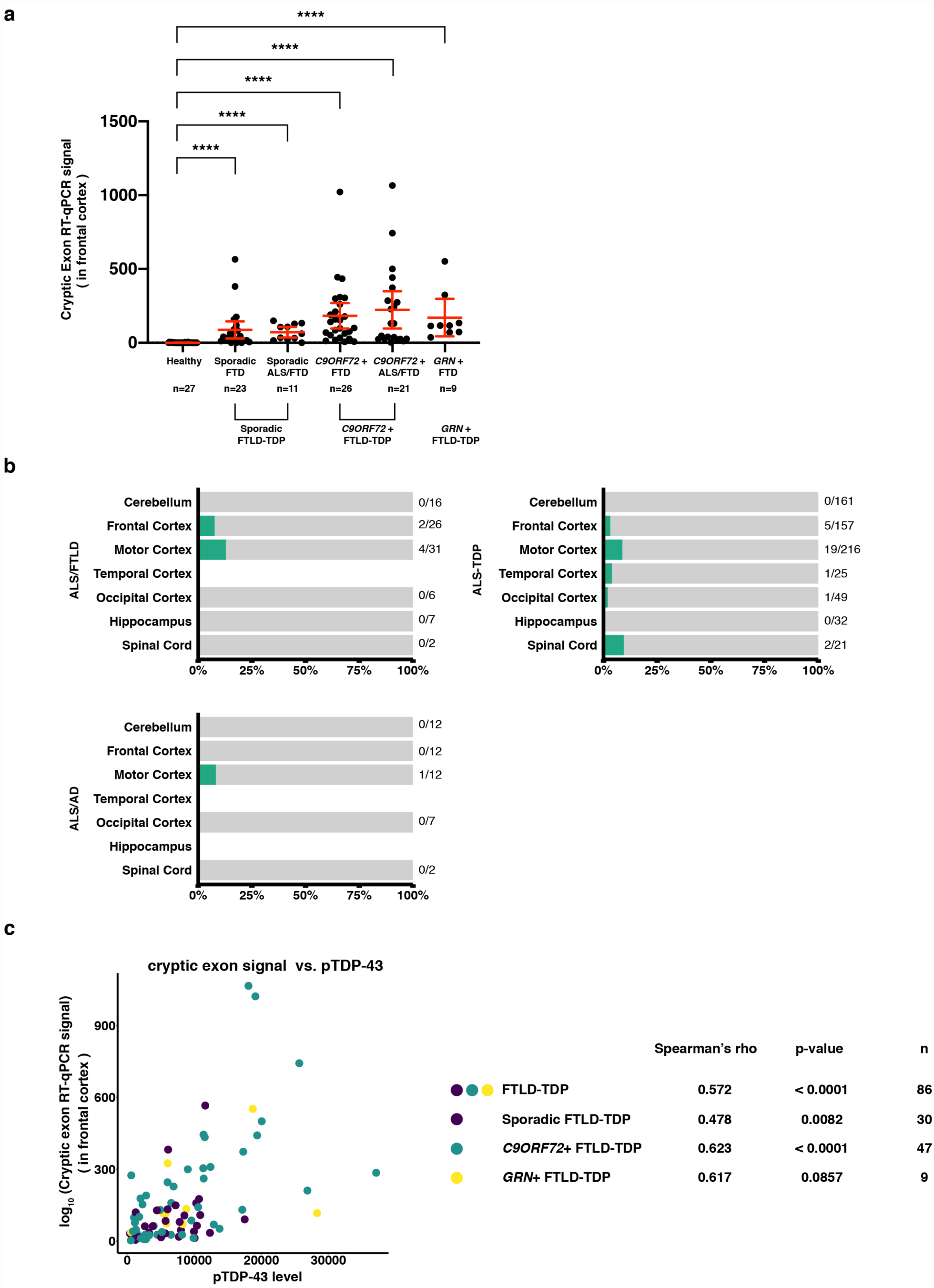
*UNC13A* cryptic exon inclusion is detected in disease relevant tissues of FTLD-TDP, ALS/FTLD, ALS-TDP and ALS/AD patients, and is correlated with phosphorylated TDP-43 levels in frontal cortices of FTLD-TDP patients. **(a)** *UNC13A* cryptic exon expression level is significantly increased in the frontal cortices of patients with FTD and FTD/ALS clinical diagnoses. *GAPDH* and *RPLP0* were used to normalize RT-qPCR (two-tailed Mann-Whitney test, ****P<0.0001; error bars represent 95% confidence intervals). **(b)** The diagnoses of these patients are not neuropathologically confirmed. Therefore, it is unclear whether TDP-43 mislocalization is present. ALS patients were categorized based on whether they carry *SOD1* mutations (ALS-SOD1 vs. ALS-TDP). ALS-AD refers to ALS patients with suspected Alzheimer’s disease. ALS-FTLD refers to patients who have concurrent FTD and ALS. **(c)** Raw data showing *UNC13A* cryptic exon signal is positively correlated with phosphorylated TDP-43 levels in frontal cortices of FTLD-TDP patients in Mayo Clinic Brain Bank (Spearman’s rho = 0.572, p-value <0.0001). Data points are colored according to patients’ reported genetic mutations. The correlation within each genetic mutation group and the corresponding p-value and sample size is also shown.

**Extended Data Fig. 6.**
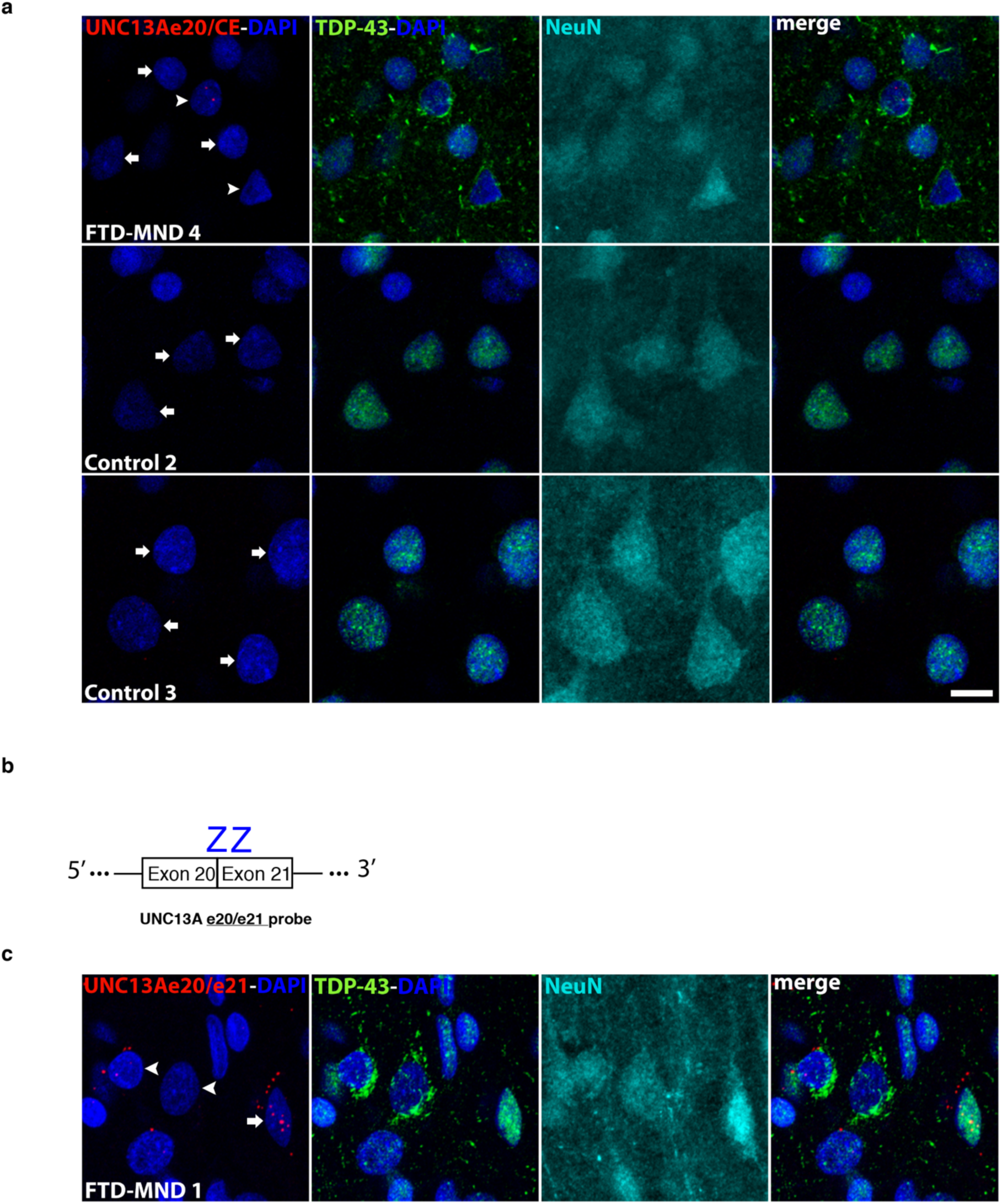
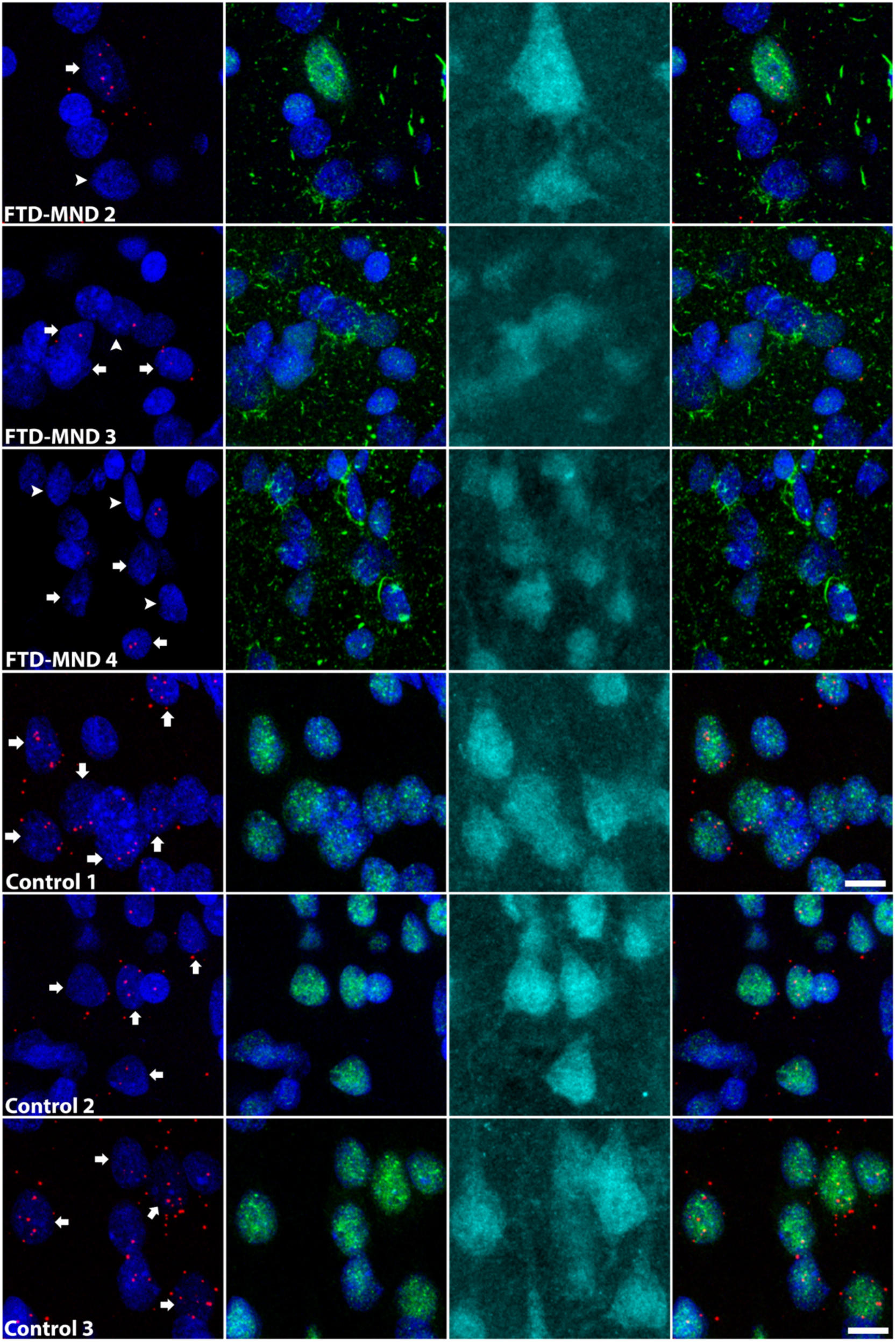
*UNC13A* cryptic splicing is associated with loss of nuclear TDP-43 in patients with FTD and motor neuron disease (MND). **(a)** Additional patients and control subjects used in the study (but not shown in Fig. 3), demonstrating *UNC13A* cryptic splicing. **(b)** The design of the *UNC13A* e20/e21 BaseScope™ probe targeting canonical *UNC13A* transcript. Each “Z” binds to the transcript independently. Both “Z”s have to be in close proximity for successful signal amplification, ensuring binding specificity. **(c)** Representative images showing expression of *UNC13A* mRNA in layer 2-3 neurons from the medial frontal pole. BaseScope™ *in situ* hybridization was used to visualize *UNC13A* mRNA, using probes that target the canonical exon20/21 junction, and combined with immunofluorescence for TDP-43 and NeuN. *UNC13A* mRNA expression is restricted to neurons (arrows) and is decreased in cells exhibiting TDP-43 proteinopathies. Arrowheads represent neurons with loss of nuclear TDP-43 and accompanying cytoplasmic inclusions, and arrows indicate neurons with normal nuclear TDP-43. Images are maximum intensity projections of a confocal image Z-stack. Scale bar equals 10 µm.

**Extended Data Fig. 7.**
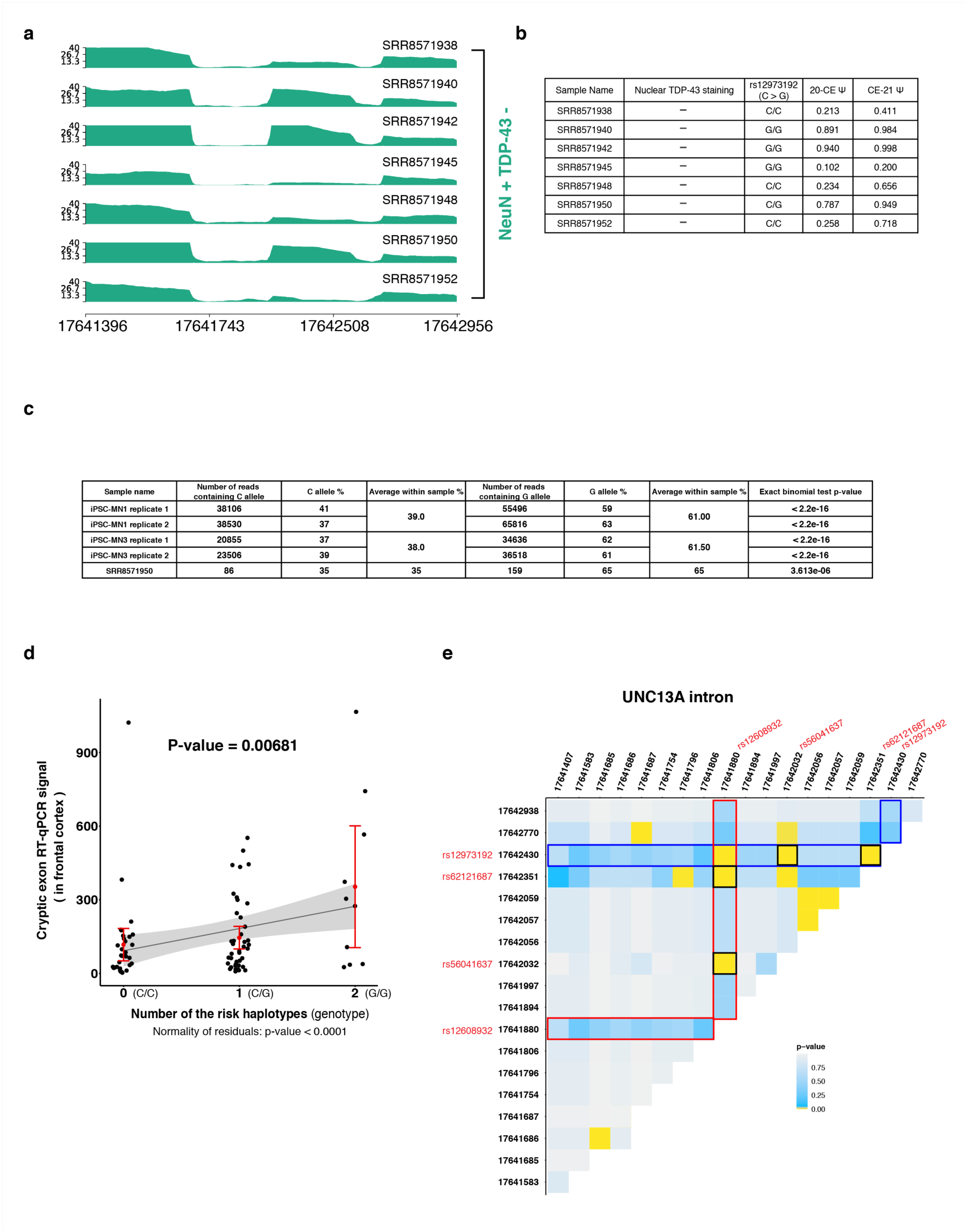
Levels of *UNC13A* cryptic exon inclusion are influenced by the number of risk haplotypes. **(a)** Visualization of RNA-Seq alignment between exon 20 and exon 21 of *UNC13A*. RNA-Seq libraries were generated from TDP-43-negative neuronal nuclei as described in Fig. 1a. **(b)** Samples that are heterozygous (C/G) or homozygous (G/G) at rs12973192 have higher relative inclusion (Ψ) of the cryptic exon with the exception of SRR8571945. **(c)** The percentages of C and G alleles in the *UNC13A* spliced variants in TDP-43 depleted iPSC-MNs and SRR8571950 neuronal nuclei. Exact binomial test was done for each replicate to test whether the observed difference in percentages differ from what was expected if both alleles are equally included in the cryptic exon. **(d)** Linear regression model using raw data show a strong correlation of 0.0186585 ^36^, we excluded it from further analysis.

**Extended Data Fig. 8.**
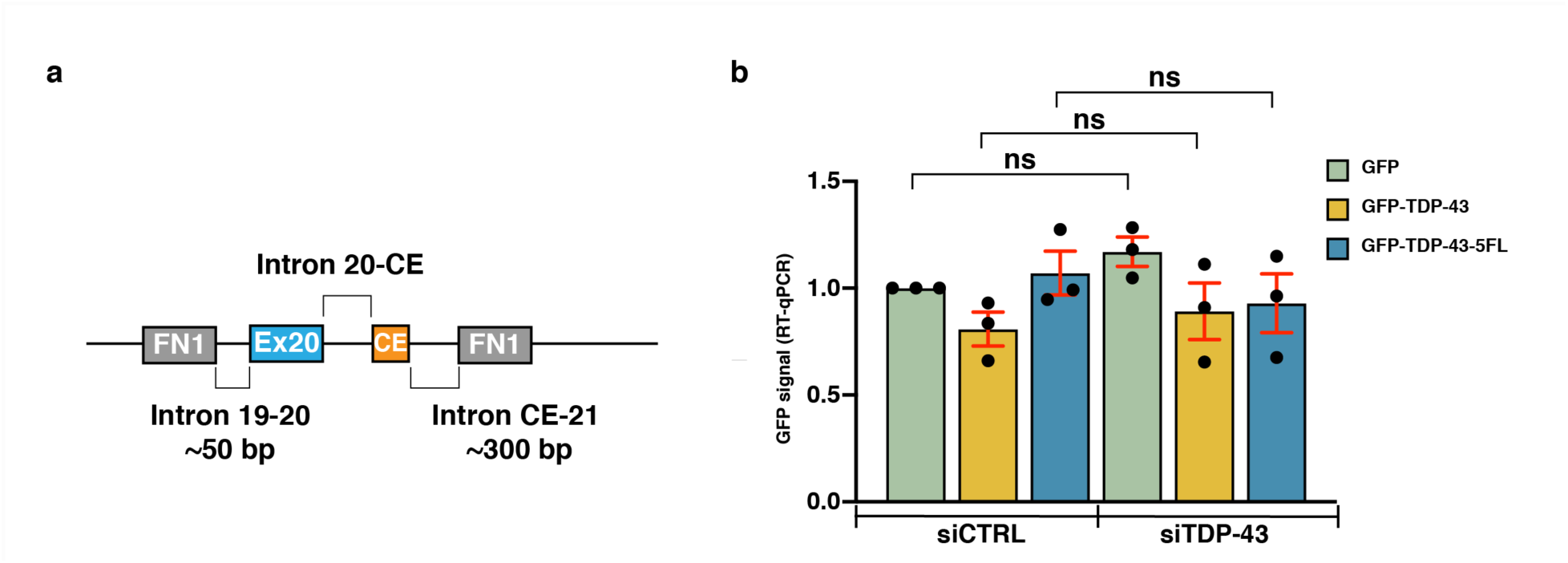
TDP-43-dependent minigene splicing reporter assay in HeLa cells. (**a**) Schematic of the pTB *UNC13A* minigene construct. The pTB *UNC13A* minigene construct containing the human *UNC13A* cryptic exon sequence and the flanking sequences upstream (50 bp at the of end of intron 19, the entire exon 20, the entire intron 20 sequence upstream of the cryptic exon) and downstream (∼300 bp intron 20) were expressed using the pTB vector, which we have previously used to study TDP-43 splicing regulation of other TDP-43 targets ^40^. (**b**) RT-qPCR of GFP demonstrating expressions of the constructs are similar across different conditions. *GAPDH* and *RPLP0* were used to normalize RT-qPCR (One-way ANOVA with Sidak’s multiple comparisons test, ns, P > 0.05).

**Extended Data Fig. 9.**
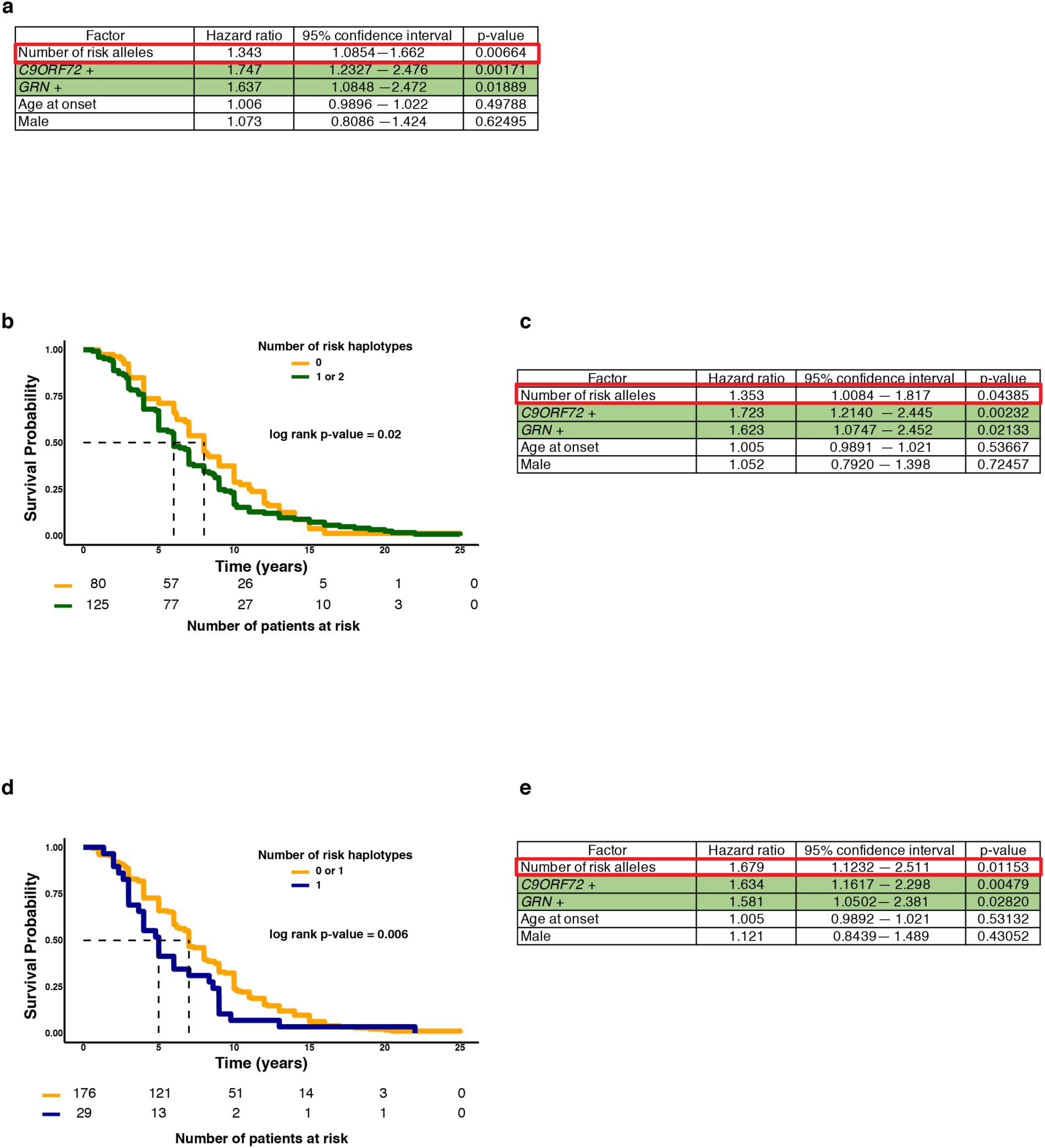
*UNC13A* risk haplotype reduces the survival time of FTLD-TDP patients. **(a)** Summary results of Cox multivariable analysis (adjusted for genetic mutations, sex and age at onset) of an additive model. **(b and d)** Survival curves of FTLD-TDP patients (n= 205, Mayo Clinic Brain Bank), according to a dominant model (b) and a 5 recessive model (d) and their corresponding risk tables. Summary results of Cox multivariable analysis (adjusted for genetic mutations, sex and age at onset) of a dominant model **(c)** and a recessive model **(e)**. Both the dominant model **(b and c)** and the recessive model **(d and e)** show that the presence of the risk haplotype can reduce the survival of FTLD-TDP patients. Dashed lines mark the median survival for each genotype. Log rank p-values were calculated using Score test. Rows colored in green indicate factors within one variable.

**Extended Data Table 1.**
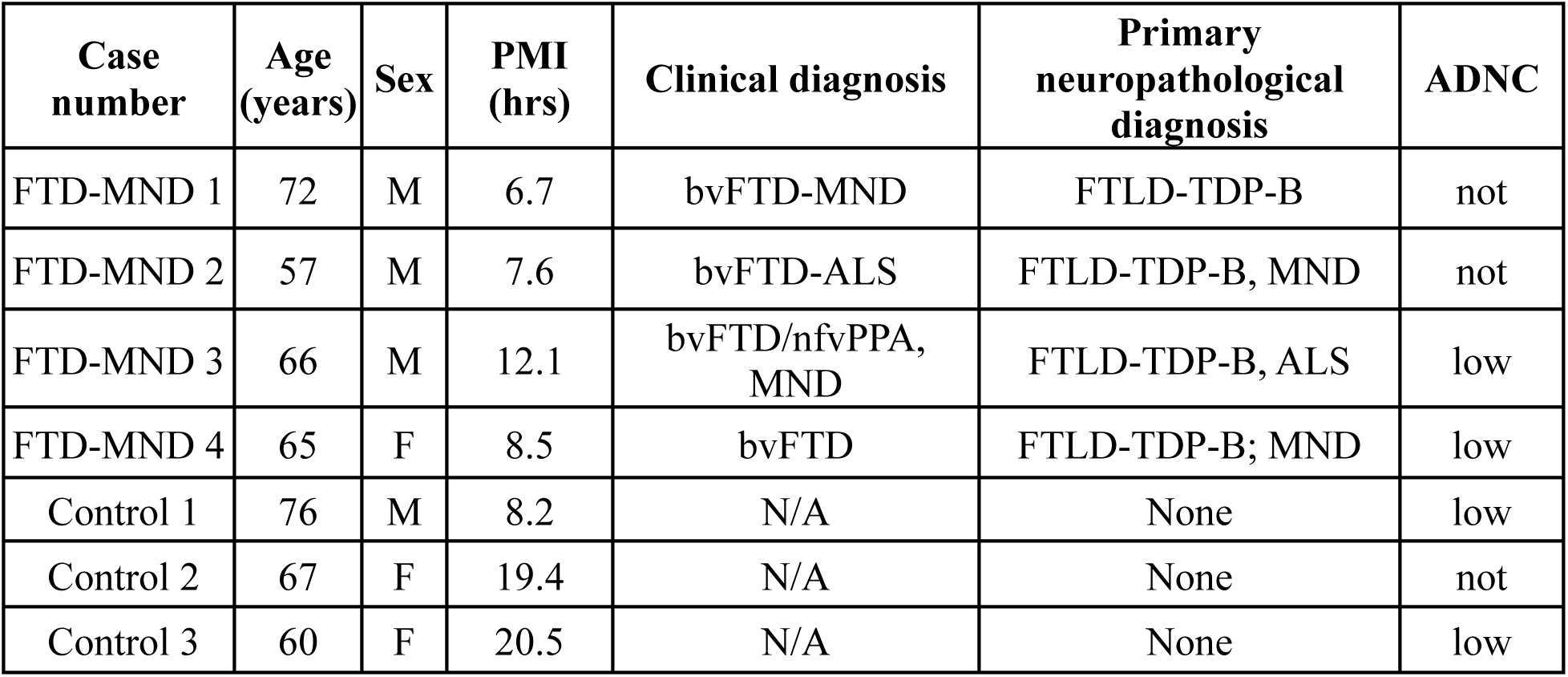
List of sporadic FTLD-TDP and control cases used for the immunohistochemical analysis of TDP-43 and *UNC13A* cryptic exon inclusion. Abbreviations: PMI – postmortem interval, bvFTD – behavioral variant frontotemporal dementia, ALS – Amyotrophic lateral sclerosis, MND – motor neuron disease, nfvPPA – non-fluent variant primary progressive aphasia, FTLD – frontotemporal lobar degeneration, ADNC - Alzheimer’s disease neuropathologic change

## Materials availability

All materials used in this study are available upon request.

## Methods

### RNA-Seq alignment and splicing analysis

Detailed pipeline v2.0.1 for RNA-Seq alignment and splicing analysis is available on https://github.com/emc2cube/Bioinformatics/sh_RNAseq.sh.

FASTQ files were downloaded from the Gene Expression Omnibus (GEO) database as GSE126543. Adaptors in FASTQ files were removed using trimmomatic (0.39) (ILLUMINACLIP:TruSeq3-PE.fa:2:30:10 LEADING:3 TRAILING:3 SLIDINGWINDOW:4:15 MINLEN:36). The quality of the resulting files was then evaluated using FastQC (v0.11.9). RNA-Seq reads were then mapped to the human (hg38) using STAR v2.7.3a.

### Splicing analysis

**MAJIQ:** Alternative splicing events were analyzed using MAJIQ (2.2) and VOILA ^12^. Briefly, uniquely mapped, junction-spanning reads were used by MAJIQ with the following parameters “majiq build -c config --min-intronic-cov 1 --simplify” to construct splice graphs for transcripts by using the UCSC transcriptome annotation (release 82) supplemented with *de novo* detected junctions. Here, *de novo* refers to junctions that were not in the UCSC transcriptome annotation, but had sufficient evidence in the RNA-Seq data (--min-intronic-cov 1). Distinct **l**ocal **s**plice **v**ariations (LSVs) were identified in gene splice graphs and the MAJIQ quantifier (majiq psi) estimated the fraction of each junction in each LSV, denoted as **p**ercent **s**pliced **i**n (PSI or Ψ), in each RNA-Seq samples. The changes in each junction’s PSI (ΔPSI or ΔΨ) between the two conditions (TDP-43-positive neuronal nuclei vs. TDP-43-negative neuronal nuclei) were calculated by using the command “majiq deltapsi”. The gene splice graphs, the posterior distribution of PSI and ΔPSI were visualized using VOILA.

**LeafCutter** (commit 249fc26 on https://github.com/davidaknowles/leafcutter): Using the already aligned RNA-Seq reads as previously described, reads that span exon-exon junction and map with a minimum of 6 np into each exon were extracted from the alignment (bam) files using filter_cs.py with the default settings. Intron clustering was performed using the default settings in leafcutter_cluster.py. Differential excision of the introns between the two conditions (TDP-43- positive neuronal nuclei vs. TDP-43-negative neuronal nuclei) were calculated using leafcutter_ds.R

### Cell culture

SH-SY5Y (ATCC) cells were grown in DMEM/F12 media supplemented with Glutamax (Thermo Scientific), 10% Fetal Bovine Serum and 10% penicillin–streptomycin at 37°C, 5% CO2. For shRNA treatments, cells were plated on Day 0, transduced with shRNA on Day 2 followed by media refresh on Day 3, and harvested for readout (RT-qPCR, immunoblotting) on Day 6. HEK293T TDP-43 knock-out cells and parent HEK-293T cells were generated as described in (37). The cells were cultured in DMEM medium (Gibco 10564011) supplemented with 10% Fetal Bovine Serum (Invitrogen 16000-044), 1% penicillin–streptomycin, 2 mM L-glutamine (Gemini Biosciences), 1x MEM non-essential amino acids solution (Gibco) at 37°C, 5% CO2.

### Immunoblotting

SH-SY5Y cells and iPSC derived motor neurons (iPSCs-MNs) were transfected and treated as above before lysis. Cells were lysed in ice-cold RIPA buffer (Sigma-Aldrich R0278) supplemented with a protease inhibitor cocktail (Thermo Fisher 78429) and phosphatase inhibitor (Thermo Fisher 78426). After pelleting lysates at maximum speed on a table-top centrifuge for 15 min at 4 °C, bicinchoninic acid (Invitrogen 23225) assays were conducted to determine protein concentrations. 60 μg (SH-SY5Y) and 30 μg (iPSCs-MNs) protein of each sample was denatured for 10 min at 70 °C in LDS sample buffer (Invitrogen NP0008) containing 2.5% 2- mercaptoethanol (Sigma-Aldrich). These samples were loaded onto 4–12% Bis–Tris gels (Thermo Fisher NP0335BOX) for gel electrophoresis, then transferred onto 0.45-μm nitrocellulose membranes (Bio-Rad 162-0115) at 100 V for 2 h using the wet transfer method (Bio-Rad Mini Trans-Blot Electrophoretic Cell 170-3930). Membranes were blocked in Odyssey Blocking Buffer (LiCOr 927-40010) for 1h then incubated overnight at room temperature in blocking buffer containing antibodies against UNC13A (1:500, Proteintech 55053-1-AP), TDP-43 (1:1,000, Abnova H00023435-M01), or GAPDH (Cell Signaling Technologies 5174S). Membranes were subsequently incubated in blocking buffer containing HRP-conjugated anti-mouse IgG (H+L) (1:2000, Fisher 62-6520) or HRP-conjugated anti-rabbit IgG (H+L) (1:2000, Life Technologies 31462) for one hour. ECL Prime kit (Invitrogen) was used for development of blots, which were imaged using ChemiDox XRS+ System (BIO-RAD). The intensity of bands was quantified using Fiji, and then normalized to the corresponding controls.

### RNA Extraction, cDNA Synthesis, and RT-qPCR/RT-PCR for detecting the *UNC13A* splice variant

Total RNA was extracted using RNeasy Micro kit (Qiagen) per manufacturer’s instructions, with lysate passed through a QIAshredder column (Qiagen) to maximize yield. RNA was quantified by Nanodrop (Thermo Scientific), with 75ng used for cDNA synthesis with SuperScript IV VILO Master Mix (Thermo Scientific). qPCR was run with 6ng cDNA input in a 20ul reaction using PowerTrack SYBR Green Master Mix (Thermo Scientific) with readout on a QuantStudio 6 Flex using standard cycling parameters (95°C for 2 minutes, 40 cycles of 95°C for 15s/60°C for 60s), followed by standard dissociation (95°C for 15s at 1.6°C/second, 60°C for 60s at 1.6°C/second, 95°C for 15s at 0.075°C/second). ΔΔCt was calculated with *RPLP0* as housekeeper and relevant shScramble as reference; measured Ct values greater than 40 were set to 40 for visualizations. The following primer pairs were used: UNC13A_CE FWD 5’-3’= TGGATGGAGAGATGGAACCT, UNC13A_CE RVS 5’-3’= GGGCTGTCTCATCGTAGTAAAC; *UNC13A* FWD 5’-3’= GGACGTGTGGTACAACCTGG, *UNC13A* RVS 5’-3’= GTGTACTGGACATGGTACGGG; TARDBP_1 FWD 5’-3’= AATTCTGCATGCCCCAGA, TARDBP_1 RVS 5’- 3’=GAAGCATCTGTCTCATCCATTTT; RPLP0_1 FWD 5’-3’= TCTACAACCCTGAAGTGCTTGAT, RPLP0_1 RVS 5’-3’ = CAATCTGCAGACAGACACTGG.

RT-PCR was conducted with 15ng cDNA input in a 100ul reaction using NEBNext Ultra II Q5 Master Mix (New England Biolabs), with resulting products visualized on a 1.5% TAE gel. The following primer pairs were used: UNC13A_19_21 FWD 5’-3’= CAACCTGGACAAGCGAACTG, UNC13A_19_21 RVS 5’-3’= GGGCTGTCTCATCGTAGTAAAC; UNC13A_CE FWD 5’-3’= TGGATGGAGAGATGGAACCT, UNC13A_CE RVS 5’-3’= GGGCTGTCTCATCGTAGTAAAC;

### shRNA cloning, lentiviral packaging, and cellular transduction

shRNA sequences originated from the Broad GPP Portal (TDP-43: AGATCTTAAGACTGGTCATTC, scramble: GATATCGCTTCTACTAGTAAG). To clone, complementary oligos were synthesized to generate 4 nt overhangs, annealed, and ligated into pRSITCH (Tet inducible U6) or pRSI16 (constitutive U6) (Cellecta). Ligations were transformed into Stbl3 chemically competent cells (Thermo Scientific) and grown at 30 °C. Large scale plasmid generation was performed using Maxiprep columns (Promega), with purified plasmid used as input for lentiviral packaging with second generation packaging plasmids psPAX2 and pMD2.G (Cellecta), transduced with Lipofectamine 2000 (Invitrogen) in Lenti-X 293T cells (Takara). Viral supernatant was collected at 48 and 72 hours post transfection and concentrated using Lenti-X Concentrator (Takara). Viral titer was established by serial dilution in relevant cell lines and readout of %BFP+ by flow cytometry, with a dilution achieving a minimum of 80% BFP+ cells selected for experiments.

### Variant validation

Variants in iPSC-derived motor neuron cells were established by PCR amplification from UNC13A exon 19 to exon 21 (UNC13A_19_21 FWD 5’-3’= CAACCTGGACAAGCGAACTG, UNC13A_19_21 RVS 5’-3’= GGGCTGTCTCATCGTAGTAAAC). Resulting products were purified using Wizard SV Gel and PCR Clean-Up columns (Promega) and submitted for Sanger and NGS (Amplicon EZ) (Genewiz).

### iPSC maintenance and differentiation into motor neurons (iPSC-MNs)

iPSC lines were obtained from public biobanks (GM25256-Corriell Institute; NDS00262, NDS00209-NINDS) and maintained in mTeSR1 media (StemCell Technologies) on matrigel (Corning). iPSCs were fed daily and split every 4-7 days using ReLeSR (StemCell Technologies) according to manufacturer’s instructions. Differentiation of iPSCs into motor neurons was carried out as previously described ^41^. Briefly, iPSCs were dissociated and placed in ultra-low adhesion flasks (Corning) to form 3D spheroids in media containing DMEMF12/Neurobasal (Thermo Fisher), N2 supplement (Thermo Fisher), and B-27 supplement-Xeno free (Thermo Fisher). Small molecules were added to induce neuronal progenitor patterning of the spheroids, (LDN193189, SB-431542, Chir99021), followed by motor neuron induction (RA, SAG, DAPT). After 14 days, neuronal spheroids were dissociated with Papain and DNAse (Worthington Biochemical) and plated on Poly-D-Lysine/Laminin coated plates in Neurobasal medium (Thermo Fisher) containing neurotrophic factors (BDNF, GDNF, CNTF; R&D Systems). For viral transductions, neuronal cultures were incubated for 18 hr with media containing lentivirus particles for shScramble, or shTDP-43. Infection efficiency of over 90% was assessed by RFP expression. Neuronal cultures were analyzed for RNA and protein 7 days post transduction.

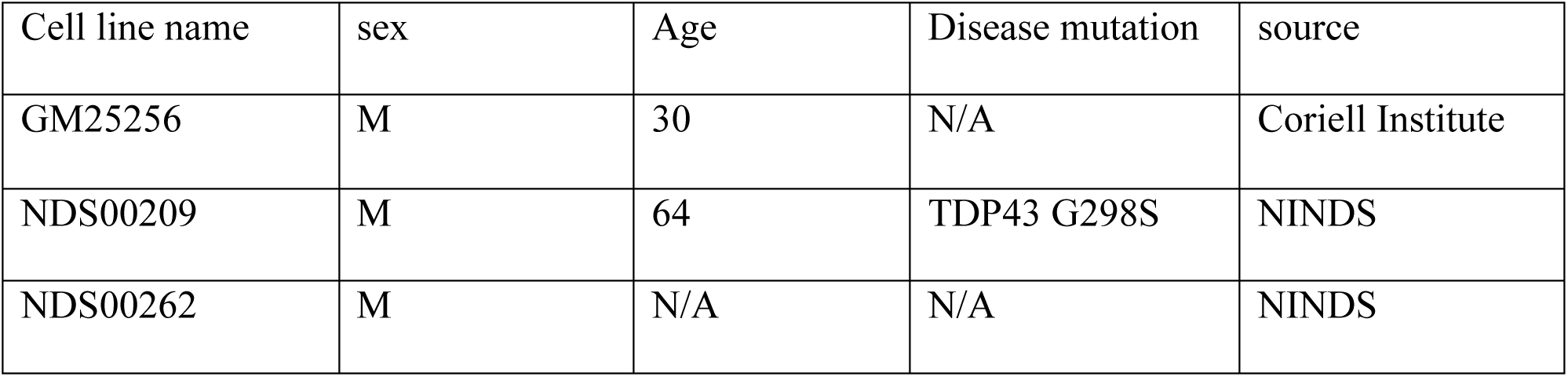

| Cell line name | sex | Age | Disease mutation | source |
| --- | --- | --- | --- | --- |
| GM25256 | M | 30 | N/A | Coriell Institute |
| NDS00209 | M | 64 | TDP43 G298S | NINDS |
| NDS00262 | M | N/A | N/A | NINDS |

### Human iPSC-neurons for detecting *UNC13A* splice variant

Complementary cDNA was available from CRISPRi-i^3^Neuron iPSCs (i^3^N) generated from our previous publication ^10^, in which TDP-43 is downregulated to about 50%. Quantitative real-time PCR (RT-qPCR) was performed using SYBR GreenER qPCR SuperMix (Invitrogen). Samples were run in triplicate, and RT-qPCRs were run on a QuantStudio™ 7 Flex Real-Time PCR System (Applied Biosystems). The following primer pairs were used: UNC13A_CE FWD 5’-3’= TGGATGGAGAGATGGAACCT, UNC13A_CE RVS 5’-3’= GGGCTGTCTCATCGTAGTAAAC. Relative quantification was determined using the ΔΔCt method and normalized to the endogenous controls RPLP0 and GAPDH (*GAPDH* FWD 5’-3’= GTTCGACAGTCAGCCGCATC, *GAPDH* RVS 5’-3’= GGAATTTGCCATGGGTGGA; RPLP0_2 FWD 5’-3’= TCTACAACCCTGAAGTGCTTGAT, RPLP0_2 RVS 5’-3’=CAATCTGCAGACAGACACTGG). We normalized relative transcript levels for wild-type *UNC13A* to that of the healthy controls (mean set to 1).

### Post-mortem brain tissues for detecting *UNC13A* splice variant

Post-mortem brain tissues from patients with FTLD-TDP and cognitively normal control individuals were obtained from the Mayo Clinic Florida Brain Bank. Diagnosis was independently ascertained by trained neurologists and neuropathologists upon neurological and pathological examinations, respectively. Written informed consent was given by all participants or authorized family members and all protocols were approved by the Mayo Clinic Institution Review Board and Ethics Committee. Complementary DNA (cDNA) obtained from 500 ng of RNA (RIN ≥ 7.0) from medial frontal cortex was available from a previous study, as well as matching pTDP-43 data from the same samples ^42^. Following standard protocols, quantitative real-time PCRs (RT-qPCR) were conducted using SYBR GreenER qPCR SuperMix (Invitrogen, Carlsbad, CA, USA) for all samples in triplicates. All primer pairs used are listed in Supplementary Table X (UNC13A_CE FWD 5’-3’= TGGATGGAGAGATGGAACCT, UNC13A_CE RVS 5’-3’= GGGCTGTCTCATCGTAGTAAAC). RT-qPCRs were run in a QuantStudio™ 7 Flex Real-Time PCR System (Applied Biosystems). Relative quantification was determined using the ΔΔCt method and normalized to the endogenous controls *RPLP0* and *GAPDH* (*GAPDH* FWD 5’-3’= GTTCGACAGTCAGCCGCATC, *GAPDH* RVS 5’-3’= GGAATTTGCCATGGGTGGA; RPLP0_2 FWD 5’-3’= TCTACAACCCTGAAGTGCTTGAT, RPLP0_2 RVS 5’-3’ 5’ CAATCTGCAGACAGACACTGG). We normalized relative transcript levels to that of the healthy controls (mean set to 1).

### Quantification of *UNC13A* splice variants

RNA-Seq data generated by NYGC ALS Consortium cohort were downloaded from the NCBI’s Gene Expression Omnibus (GEO) database (GSE137810, GSE124439, GSE116622, and GSE153960). We used the 1658 available and quality-controlled samples classified as described in ^10^. After pre-processing and aligning the reads to human (hg38) as described previously, we estimated the expression of the full-length *UNC13A* using RSEM (v1.3.2). The average TPM of UNC13A across all the tissue samples from all the individuals was 10.5 on average. PCR duplicates were removed using MarkDuplicates from Picard Tools (2.23.0) using the command “MarkDuplicates REMOVE_DUPLICATES=true CREATE_INDEX=true”. Reads that span either “Exon 19-Exon 20” junction, “Exon 20-CE” junction, “CE- Exon 21” junction, or “Exon 20-exon 21” junction were quantified using bedtools (2.27.1) using the command “bedtools intersect -split”. Because of the relatively low level of expression of *UNC13A* in post-mortem tissues and the heterogeneity of the tissues, it is possible that not all tissues have enough detectable *UNC13A* for us to detect the splice variants. Since *UNC13A* contains more than 40 exons and RNA-Seq coverages of mRNA transcripts are often not uniformly distributed ^43^, we looked at reads spanning “Exon 19-Exon 20” junction, which is included in both the canonical isoform and the splice variant, and there is a strong correlation (Pearson’s r = 0.99) between the numbers of reads mapped to “Exon 19- Exon 20” junction and “Exon 20-Exon 21” junction. We observed that samples that have at least 2 reads spanning either “Exon 20-CE” junction or “CE-Exon 21” junction have at least either *UNC13A* TPM = 1.55 or 20 reads spanning “Exon 19- Exon 20” junction. Therefore, we selected the 1151 samples that had a TPM ⩾ 1.55, or at least 20 reads mapped to the “Exon 19-Exon 20” junction as samples suitable for *UNC13A* splice variant analysis.

### Determination of rs12608932 and rs12973192 SNP genotype in human postmortem brain

Genomic DNA (gDNA) was extracted from human frontal cortex using Wizard Genomic DNA Purification Kit (Promega), according to the manufacturer’s instructions. TaqMan SNP genotyping assays were performed on 20 ng of gDNA per assay, using a commercial pre-mixture consisted of a primer pair and VIC/FAM labeled probes specific for each SNP (Cat#4351379, assay ID “43881386_10” for rs12608932 and “11514504_10” for rs12973192, Thermo Fisher Scientific), and run on a QuantStudio™ 7 Flex Real-Time PCR system (Applied Biosystems), according to the manufacturer’s instructions. The PCR-programs were 60°C for 30 s, 95°C for 10min, 40 cycles of 95°C for 15s and, 60°C (rs12973192) or 62.5°C for 1min (rs12608932), and 60°C for 30s.

### Splicing Reporter Assay

Minigene constructs were designed in silico, synthesized by GeneScript and sub-cloned into a vector with the GFP splicing control. HEK293T TDP-43 knock-out cells and the parent HEK- 293T cells were seeded into standard P12 tissue culture plates (at 1.6 × 10^5^ cells/well), allowed to adhere overnight and transfected with the indicated splicing reporter constructs (400 ng/well) using Lipofectamine 3000 Transfection Reagent (Invitrogen). Each reporter comprised one of the splicing modules (shown in Fig. 4E), which is expressed from a bidirectional promoter. Twenty-four hours after transfection, RNA was extracted from these cells using PureLink RNA Mini Kit (Life Technologies) according to the manufacturer’s protocol, with on-column PureLink DNase treatment. The RNA was reverse transcribed into cDNA using the High Capacity cDNA Reverse Transcription Kit (Invitrogen) according to the manufacturers’ instructions. PCRs were performed using OneTaq 2X Master Mix with Standard Buffer (NEB) using the following primers: mCherry FWD 5’-3’= GTTCATGCGCTTCAAGGTG, mCherry RVS 5’-3’=TTGGTCACCTTCAGCTTGG; EGFP FWD 5’-3’=ACAGGTACTGTGCCTATCAAAG; EGFP RVS 5’-3’= TGTGGCGGATCTTGAAGTTAG on a Mastercycler Pro (Eppendorf) thermocycler PCR machine. PCR products were separated by electrophoresis on a 1.5% TAE gel and imaged ChemiDox XRS+ System (BIO-RAD).

### Generation of pTB *UNC13A* minigene construct

The pTB *UNC13A* minigene construct containing the human *UNC13A* cryptic exon sequence and the nucleotide flanking sequences upstream (50 bp at the of end of intron 19, the entire exon 20, the entire intron 20 sequence upstream of the cryptic exon) and downstream (∼300 bp intron 20) of the cryptic exon were amplified from human genomic DNA using the following primers: FWD 5’-3’=AGGTCATATGCACTGCTATAGTGGGAAGTTC and RVS 5’-3’=CTTACATATGTAATAACTCAACCACACTTCCATC; and subcloned into the NdeI site of the pTB vector. Note we have previously used a similar approach to study TDP-43 splicing regulation of other TDP-43 targets ^44^.

### Rescue of *UNC13A* splicing using the pTB minigene and TDP-43 overexpression constructs

HeLa cells were grown in Opti-MEM I Reduced Serum Medium, GlutaMAX Supplement (Gibco) plus 10% fetal bovine serum (Sigma) and 1% penicillin/streptomycin (Gibco). For double-transfection and knockdown experiments, cells were first transfected with 1.0 µg of pTB *UNC13A* minigene construct and 1.0 µg of one of the following plasmids: GFP, GFP-TDP-43 or GFP-TDP- 43 5FL (constructs to express GFP-tagged TDP-43 proteins have been previously described ^40, 44^, in serum-free media and using Lipofectamine 2000 following manufacturer’s instructions (Invitrogen). Four hours following transfection, media was replaced with complete media containing siLentfect (Bio-Rad) and siRNA complexes (AllStars Neg. Control siRNA or siRNA against *TARDBP* 3’UTR, a region not included in the TDP-43 overexpression constructs) (Qiagen) following the manufacturer’s protocol. Cycloheximide (Sigma) was added at a final concentration of 100 µg/ml at six hours prior harvesting the cells. Then cells were harvested and RNA extracted using TRIzol Reagent (Zymo Research), following manufacturer’s instructions. Approximately 1µg of RNA was converted into cDNA using the High Capacity cDNA Reverse Transcription Kit with RNA inhibitor (Applied Biosystems). The RT-qPCR assay was performed on cDNA (diluted 1:40) with SYBR GreenER qPCR SuperMix (Invitrogen) using QuantStudio7™ Flex Real-Time PCR System (Applied Biosystems). All samples were analyzed in triplicates. The RT-qPCR program was as follows: 50°C for 2 min, 95°C for 10 min, and 40 cycles of 95 °C for 15 s and 60°C for 1 min. For dissociation curves, a dissociation stage of 95°C for 15 s, 60°C for 1 min and 95°C for 15 s was added at the end of the program. Relative quantification was determined using the ΔΔCt method and normalized to the endogenous controls *RPLP0* and *GAPDH*. We normalized relative transcript levels for wild-type *UNC13A* and GFP to that of the control siRNA condition (mean set to 1).

The following primer pairs were used: UNC13A_CE_minigene FWD 5’-3’= GATTGAACAGATGAATGAGTGATGA, UNC13A_CE_minigene RVS 5’-3’ = TGTCTGGACCAATGTTGGTG; GFP_OE FWD 5’-3’= GAAGCGCGATCACATGGT, GFP_OE RVS 5’-3’ =CCATGCCGAGAGTGATCC; *GAPDH* FWD 5’-3’=GTTCGACAGTCAGCCGCATC, *GAPDH* RVS 5’-3’ = GGAATTTGCCATGGGTGGA; RPLP0_2 FWD 5’-3’= TCTACAACCCTGAAGTGCTTGAT, RPLP0_2 RVS 5’-3’ = CAATCTGCAGACAGACACTGG; TARDBP_2 FWD 5’-3’= TGGACGATGGTGTGACTGCAA, TARDBP_2 RVS 5’-3’ = AGAGAAGAACTCCCGCAGCTCA.

### In situ hybridization *UNC13A* cryptic exon analysis in postmortem brain samples

#### Patients and diagnostic neuropathological assessment

Postmortem brain tissue samples used for this study were obtained from the University of California San Francisco (UCSF) Neurodegenerative Disease Brain Bank. Supplemental Table 1 provides demographic, clinical, and neuropathological information. Consent for brain donation was obtained from subjects or their surrogate decision makers in accordance to the Declaration of Helsinki, and following a procedure approved by the UCSF Committee on Human Research. Brains were cut fresh into 1 cm thick coronal slabs and alternate slices were fixed in 10% neutral buffered formalin for 72 h. Blocks from medial frontal pole were dissected from the fixed coronal slabs, cryoprotected in graded sucrose solutions, frozen, and cut into 50 µm thick sections as described previously ^45^. Clinical and neuropathological diagnosis were performed as described previously (*44*). Subjects were selected based on clinical and neuropathological assessment. Patients selected had a primary clinical diagnosis of behavioral variant frontotemporal dementia (bvFTD) with or without amyotrophic lateral sclerosis (ALS)/motor neuron disease (MND) and 2) a neuropathological diagnosis of frontotemporal lobar degeneration (FTLD)-TDP, Type B. We excluded subjects if they had a known disease-causing mutation, post-mortem interval ≥ 24 h, Alzheimer’s disease neuropathologic change > low, Thal amyloid phase > 2, Braak neurofibrillary tangle stage > 4, CERAD neuritic plaque density > sparse, and Lewy body disease > brainstem predominant ^45^.

#### *In situ* hybridization (ISH) and immunofluorescence

To detect single RNA molecules, a BaseScope Red Assay kit (ACDBIO, USA) was used. One 50 µm thick fixed frozen tissue section from each subject was used for staining. Experiments were performed under RNase free conditions as appropriate. Probes that target the transcript of interest, UNC13A, specific to either the mRNA (exon20/21 junction) or the cryptic exon containing spliced target (exon20/cryptic exon junction) were used. Positive (Homo sapiens PPIB) and negative (Escherichia coli DapB) control probes were also included. *In situ* hybridization was performed based on vendor specifications for the BaseScope Red Assay kit. Briefly, frozen tissue sections were washed in PBS and placed under an LED grow light (HTG Supply, LED-6B240) chamber for 48 h at 4 °C to quench tissue autofluorescence. Sections were quickly rinsed in PBS and blocked for endogenous peroxidase activity. Sections were transferred on to slides and dried overnight. Slides were subjected to target retrieval and protease treatment and advanced to ISH. Probes were detected with TSA Plus-Cy3 (Akoya Biosciences) and subjected to immunofluorescence staining with antibodies to TDP-43 (rabbit polyclonal, Proteintech, RRID: AB_615042) and NeuN (Guinea pig polyclonal, Synaptic systems) and counterstained with DAPI (Life Technologies) for nuclei.

#### Image acquisition and analysis

Z-stack images were captured using a Leica SP8 confocal microscope with an 63x oil immersion objective (1.4 NA). For RNA probes, image capture settings were established during initial acquisition based on PPIB and DAPB signal and remained constant across *UNC13A* probes and subjects. TDP-43 and NeuN image capture settings were modified based on staining intensity differences between cases. For each case, 6 non-overlapping Z-stack images were captured across cortical layers 2-3. RNA puncta for the UNC13A cryptic exon were quantified using the “analyze particle” plugin in ImageJ. Briefly, all images were adjusted for brightness using similar parameters and converted to maximum intensity Z-projections, images were adjusted for auto-threshold (intermodes), and puncta were counted (size: 6-infinity, circularity – 0-1).

### Linkage Disequilibrium analysis

Recalibrated VCF files generated by GATK HaplotypeCallers were downloaded from Answer ALS in July 2020. VCFtools (0.1.16) were used to filter for sites that are in intron 20-21. The filtered VCF files were merged using BCFtools (1.8). Since there are sites that contain more than 2 alleles, we tested for genotype independence using the chi-squared statistics by using the command “vcftools --geno-chisq --min-alleles 2 --max-alleles 8” (4.0.0).

### Statistical methods

Survival curves were compared using the coxph function in the survival (3.1.12) R package, which fits a multivariable Cox proportional hazards model that contains sex, reported genetic mutations and age at onset, and performs a Score (log-rank) test. Effect sizes are reported as the hazard ratios. Proportional Hazards assumptions were tested using cox.zph() function. The survival curves were plotted using ggsurvplot() in suvminer (v.0.4.8) R package.

Correlations between the cryptic exon signal and phosphorylation levels of TDP-43 or number of risk haplotypes were done after filtering out all the samples that do not have the cryptic exon signal (n = 4). Linear mixed effects models were analyzed using lmerTest R package (3.1.3).

Statistical analyses were performed using R (version 4.0.0), or Prism 8 (GraphPad), which were also used to generate graphs.

